# Antiviral defence is a conserved function of diverse DNA glycosylases

**DOI:** 10.1101/2025.10.29.685425

**Authors:** Landon J. Getz, Amy L. Qian, Y. Vivian Liu, Sam R. Fairburn, Mahnoor S. Butt, Yan-Jiun Lee, Peter R. Weigele, Karen L. Maxwell

## Abstract

Bacteria are frequently attacked by viruses, known as phages, and rely on diverse defence systems like restriction endonucleases and CRISPR-Cas to survive. While phages can evade these defences by covalently modifying their DNA, these non-canonical nucleobases create a strong selective pressure for host proteins that can recognize and exploit them. Here, using a structure-guided discovery approach, we identify widespread families of DNA glycosylases that protect bacteria against phages that incorporate modified guanine bases into their DNA. Despite high sequence variation, these enzymes share a conserved glycosylase fold and occur across bacterial lineages. We also uncover a distinct glycosylase superfamily that defends against phages with thymidine modifications, showing that glycosylases have repeatedly evolved as antiviral defences. Together, these findings reveal DNA glycosylases as versatile effectors of bacterial immunity and underscore structure-guided discovery as a powerful strategy for uncovering hidden layers of antiviral defence.

## Introduction

Bacteria use a variety of DNA-targeting systems to defend against viral predation, including restriction enzymes, CRISPR–Cas systems, and Argonaute nucleases (*1–4*). These effectors recognize and degrade foreign nucleic acids with high specificity and serve as central components of the prokaryotic immune arsenal. Recent efforts to systematically catalogue bacterial defences have uncovered a broad diversity of nucleases, including enzymes with novel folds, catalytic mechanisms, and modes of regulation (*5–9*). These discoveries highlight how selective pressure from phage infection has driven functional diversification of nuclease-based immunity.

To evade these defences, many phages synthesize and incorporate chemically modified bases such as 7-deazaguanine (*10*, *11*) or 5-hydroxymethylcytosine (*12–14*) into their DNA. These modifications shield phage DNA during infection yet also create potential signatures of non-self that can be recognized by bacteria. Type IV restriction enzymes exploit this by cleaving modified bases (*15–17*), and a recent study revealed Brig1, a uracil DNA glycosylase (UDG) homologue, as an unexpected phage defence enzyme (*18*). Glycosylases comprise a large and structurally diverse enzyme family (table S1) best known for their role in base excision repair, where they cleave the N-glycosidic bond between a damaged or chemically modified base and the deoxyribose sugar, creating an abasic site that is subsequently processed by repair enzymes (*19*, *20*). Brig1 is striking because instead of maintaining genome integrity through DNA repair, it excises glycosylated cytosines from T-even phages and triggers viral genome destruction. This discovery revealed that a housekeeping repair glycosylase can be repurposed for antiviral defence and suggested that other DNA glycosylases may likewise serve as hidden components of bacterial immunity.

## Results

In a previous screen for novel defence systems in *Vibrio parahaemolyticus*, nine new systems were discovered (*21*). Strikingly, two of these were unannotated proteins whose predicted structures resembled members of the helix-hairpin-helix (HhH) glycosylase superfamily (Fig. 1A). The fact that two of nine newly discovered systems both converged on glycosylase-like folds was unexpected and pointed to glycosylases as a potentially widespread but underappreciated theme in antiviral immunity. Protein–DNA modelling using AlphaFold3 supported a role for both proteins in base-modification sensing as 7-deazagunanine modified nucleotides were positioned in conserved binding pockets in both Dag1 and Dag2, while no interaction was predicted with unmodified DNA. In the DNA bound models, the modified base flipped out from the duplex into a pocket on the protein surface (Fig. 1B), a hallmark of glycosylase mechanism (*19*) that suggests Dag1 and Dag2 engage DNA in a manner consistent with base excision activity.

**Fig 1.**
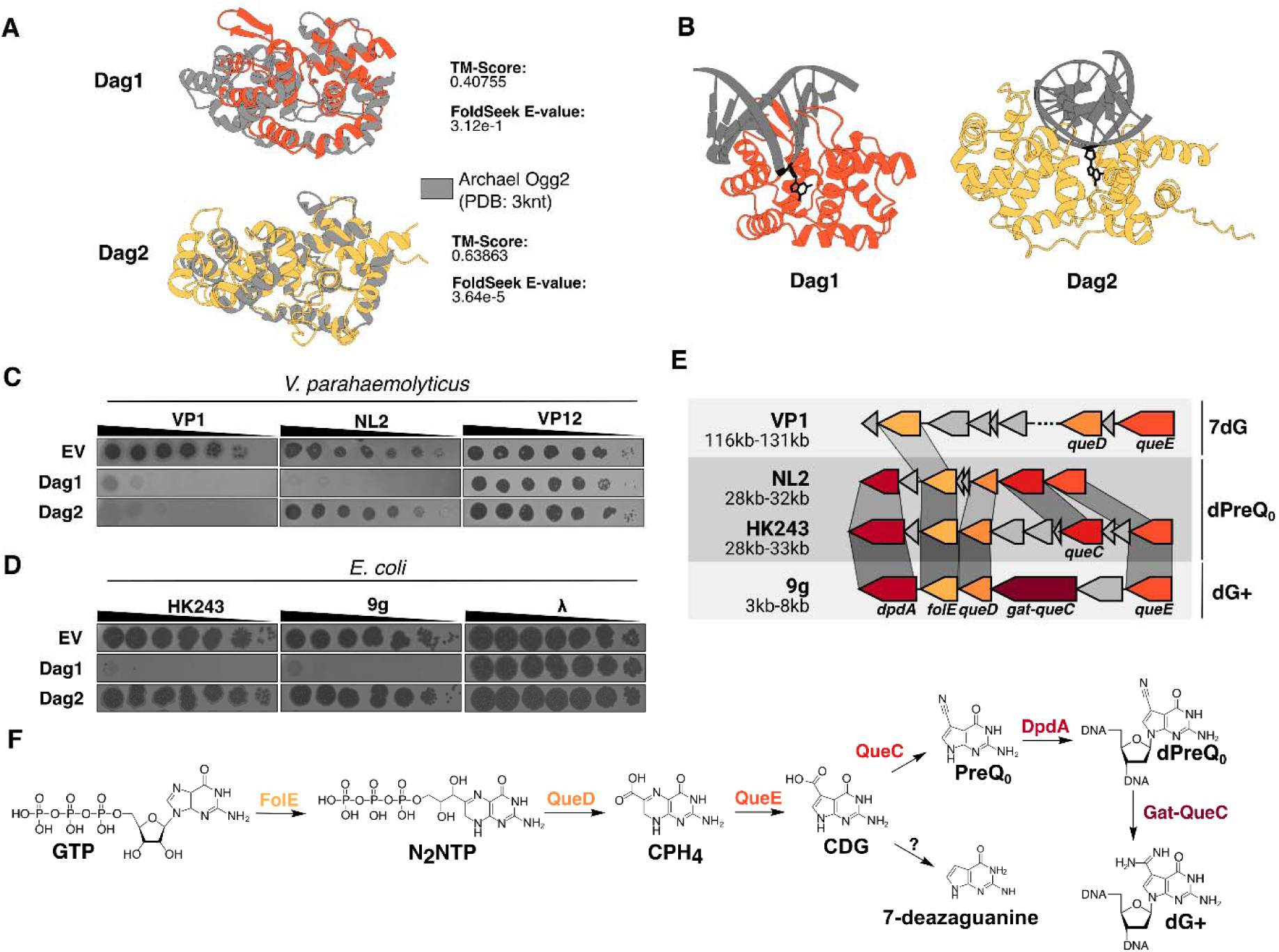
Dagl and Dag2 are structurally similar to DNA glycosylases and specifically target G-modified phages. **(A)** FoldSeek was used to identify structurally related proteins to Dagl (orange) and Dag2 (yellow). A FoldSeek identified structure is shown overlaid with Dagl and Dag2 in grey and associated TM-Scores and E-values are shown. **(B)** AlphaFold3 models of Dagl and Dag2 were generated along with double­ stranded DNA containing a 7-deazaguanine DNA base (black; Dagl ipTM: 0.58, Dag2 ipTM: 0.56). Phage plating assays in both *V parahaemolyticus* **(C)** and *E. coli* **(D)** were performed against Dagl and Dag2. Images shown are representative of three biological replicates. **(E)** Clinker was used to generate genome alignments of VP1, NL2, HK243, and 9g. Genes involved in modifying phage genomic DNA are coloured and labelled. Grey links between the genes indicate level of amino acid identity (darker grey = higher amino acid identity). **(F)** Schematic diagram of the GTP modification pathway found in the phages shown. Proteins involved are labelled if known.

### Dag1 and Dag2 provide defence against phages with modified guanines

To investigate the defence activity of Dag1 and Dag2, we expressed them from a plasmid in *V. parahaemolyticus, Escherichia coli* and *Pseudomonas aeruginosa*, and assessed their ability to inhibit replication of a panel of diverse phages (table S2). Dag1 blocked replication of *Vibrio* phages VP1 and NL2, *E. coli* phages 9g and HK243, and *P. aeruginosa* phage PHP2, while Dag2 inhibited only phage VP1 (Fig. 1C, D and fig. S1), indicating that they provide selective defence rather than broad antiviral activity. Previous work showed that *E. coli* phage 9g incorporates deoxyarchaeosine (dG), a modified guanine base originally identified in archaeal tRNA (*22*), suggesting that Dag1 may recognize guanine base modifications as a marker of phage infection. Modified guanine targeting was supported by the observation that neither predicted glycosylase affected phages with unmodified genomes such as λ or VP12, nor those with unrelated modifications including β-glucosyl-5-hydroxymethyl-cytosine (5-gmC) in phage T4, 5-hydroxymethylcytosine (5-hmC) in T4*gt*, or N^6^-methylcarbamoyladenine (6-NcmdA) in phage Mu (Fig. 1C, D and fig. S1)

To determine whether targeted phages contained DNA base modifications, we purified DNA from HK243, NL2, and VP1 and analyzed them by high-performance liquid chromatography-tandem mass spectrometry (HPLC-MS) (fig. S2A). HK243 and NL2 contained the modified base preQ_0_, representing ∼30% of total guanine incorporation as estimated from normalized dG:dC peak areas (fig. S2A,B, and Table 1). VP1 carried a distinct variant, 7-deazaguanine, with HPLC-MS revealing a complete absence of canonical guanine, consistent with full substitution (fig. S2A and C). Such total replacement is rare but has been reported in Firehammerviruses that incorporate 2′-deoxy-7-deaza-(aminoethyl)-guanine (dADG) (*23*). As expected, VP12 and λ genomes contained only canonical nucleotides (fig. S2A). Together, these data support a model in which Dag1 and Dag2 act as defence-associated glycosylases that recognize guanine base modifications as signatures of viral infection.

**Table 1.**
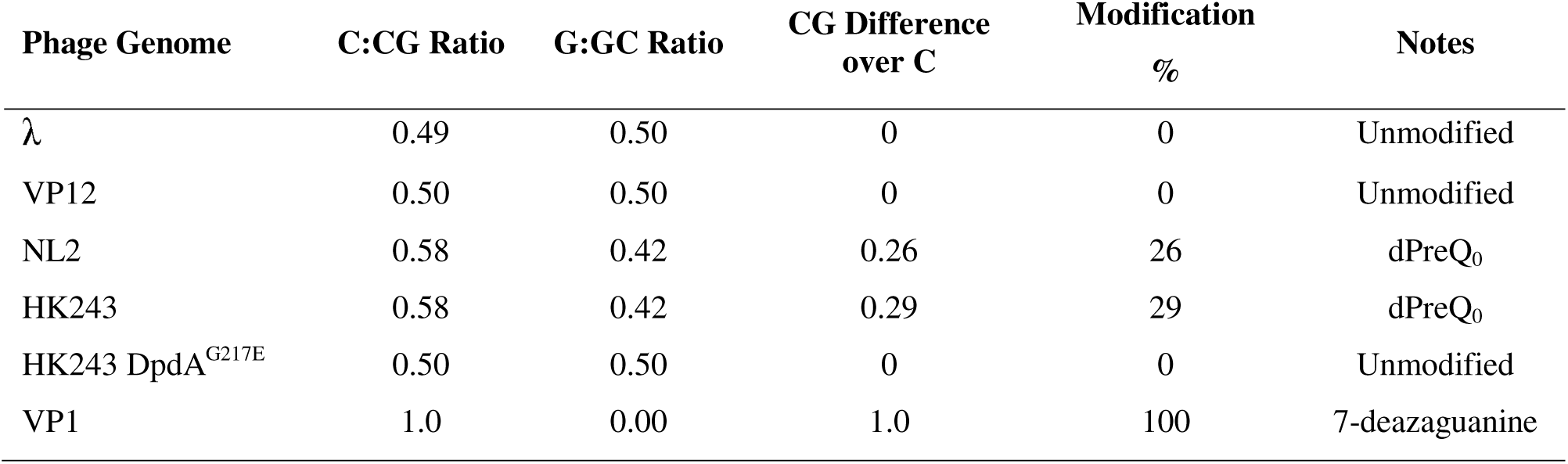
Phage Genome Modifications Detected by HPLC-MS.

Consistent with these findings, genome analysis of all five targeted phages revealed conserved pathways for guanine modification. Each phage encodes FolE, a GTP cyclohydrolase I that catalyzes the first step in the queuosine/archaeosine (Q/G) biosynthesis pathway (Fig. 1E and F). This pathway, normally used for tRNA modification in bacteria and archaea, has been adapted by phages to incorporate 7-deazaguanine derivatives into their DNA and thereby evade host restriction enzymes (*10*). We next analyzed these genomes using Domainator (*24*), a hidden Markov model–based tool that identifies enzymatic domains even in highly diverged sequences, allowing detection of pathway components that simple sequence searches might miss. The annotation revealed additional components of guanine modification pathways in NL2, HK243, and PHP2, including *queC*, *queD*, and *dpdA*, which mediate synthesis and incorporation of preQ_0_, a 7-cyano-7-deazaguanine base (Fig. 1E and F). In VP1, distant homologues of *folE*, *queC*, and *queD* were detected, further confirming incorporation of 7-deazaguanine rather than preQ_0_ (Fig. 1E and F).

We next asked whether Dag1, which showed the broadest defence profile, could directly cleave modified phage genomes *in vitro*. Purified Dag1 (fig. S3A) was incubated with phage DNA, and integrity was assessed by agarose gel electrophoresis. DNA from VP1, 9g, and HK243, which all carry modified guanine bases, was degraded in a Dag1 concentration-dependent manner (Fig. 2A). By contrast, unmodified or 5-gmC-modified DNA from phage λ and T4, respectively, remained intact. These results demonstrate that Dag1 selectively cleaves phage DNA containing guanine modifications while leaving canonical DNA or genomes bearing unrelated base modifications intact.

**Fig. 2.**
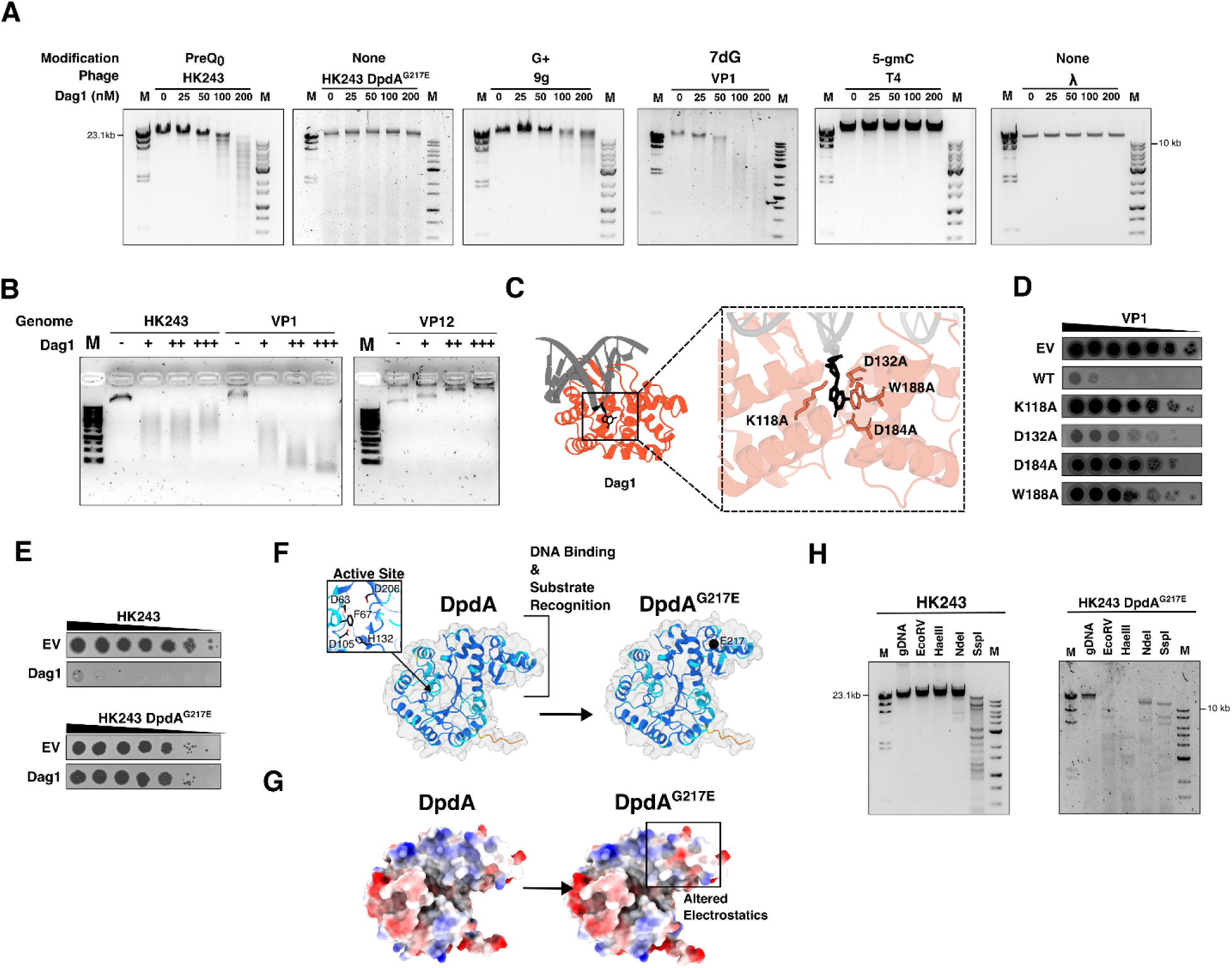
Dagl selectively degrades restriction-resistant guanine modified phage DNA. **(A)** Purified phage DNA was incubated in the presence of increasing concentrations (nM) of Dagl for 30 minutes at 37°C, followed by agarose gel electrophoresis. Nucleic acid was visualized using SYBR Safe. Left M - λ Ladder, Right M - lOkb Ladder. Maximum sizes of the ladders are indicated. Known DNA modification is shown. **(B)** *In vitro* Dagl DNA digestion experiments were performed on HK243, VP1, and VP 12 DNA with increasing amounts of Dagl enzyme. 1% agarose gel electrophoresis was performed at 4°C and 20V to prevent heat-induced degradation of abasic DNA. **(C)** Dagl AlphaFold3 model (orange) with 7-deazaguanine (black). Key interacting active site residues are shown and labelled. **(D)** Site-directed mutagenesis was performed to mutate the individual active site residues to alanine, followed by expression and phage plating assays. Phage plating experiments were performed with Dagl targeted phage VP1 in *V. parahaemolyticus.* Images are representative of three biological replicates. **(E)** Phage plating assays were performed following the isolation of a Dagl escape mutant (HK243 DpdA^G217E^) on cells containing either empty vector or those expressing Dagl from a plasmid. Images are representative of three biological replicates. (F) Cartoon schematic of the AlphaFold3 predicted DpdA protein from HK243. The putative active sites are shown. Cartoons are coloured by pLDDT, using AlphaFold3 colouring (dark blue, pLDDT >90; light blue, 9O>pLDDT>7O; yellow, low 7O>pLDDT>5O; orange, very low pLDDT<5O). **(G)** Coulombic electrostatic potential of the same proteins as in **(F).** were calculated using ChimeraX and are shown for each residue (Blue = +10, Red = -10). Altered electrostatics at E217 in the mutant are indicated by a black box. **(H)** Restriction analysis of HK243 and HK243 DpdA^G217E^. Restriction enzymes are indicated. Ladders (M) are same as in (A). 1.0% agarose gels are shown, and nucleic acid was visualized using SYBR Safe.

DNA glycosylases can be broadly categorized as monofunctional or bifunctional enzymes (*19*). Monofunctional glycosylases generate an abasic site that is processed by separate endonucleases, whereas bifunctional enzymes possess intrinsic lyase activity that also nicks the DNA strand. To determine whether Dag1 is mono- or bifunctional, we repeated the *in vitro* DNA degradation assays under conditions that prevent non-enzymatic strand cleavage at abasic sites (*18*, *25*). The HK243 genome, which contains ∼30% modified guanines, was fragmented into broad DNA smears, while the fully modified VP1 genome was degraded to very small fragments, showing that cleavage extent scales with modification density (Fig. 2B). The fragmentation was observed even under electrophoresis conditions that minimized heat and electrolysis (4 °C, low voltage), suggesting that strand breakage was enzyme-driven rather than the result of abasic site instability. As expected, unmodified VP12 DNA remained intact. Together, these results imply that Dag1 acts as a bifunctional glycosylase capable of both excising modified bases and cleaving the DNA backbone.

The structural model of Dag1 bound to a 7-deazaguanine–modified DNA duplex using AlphaFold3 revealed a compact pocket oriented toward the DNA, with several residues positioned to contact the modified base (Fig. 2C). The model identified one lysine (K118) and two aspartates (D132 and D184) near the modified base; each positioned such that they could participate in catalysis. An additional residue, W188, was located adjacent to the modified base, where it likely stabilizes it through stacking interactions. These four residues define the core of the predicted active site and provided a basis for testing their roles in substrate recognition and catalysis. To assess their functional importance, we generated alanine mutants and measured defence activity against phage VP1 (Fig. 2D). The K118A and D184A substitutions completely abolished defence, consistent with an essential role in catalysis. W188A likewise abolished activity, consistent with its predicted role in base stabilization. By contrast, D132A retained partial activity: plaques on Dag1-expressing cells were turbid and the efficiency of plating decreased by about 10-fold relative to wild type, suggesting that this residue contributes less to glycosylase activity. Western blotting confirmed that all variants were expressed and soluble at levels comparable to wild-type Dag1 (fig. S3B and C), demonstrating that the observed phenotypes reflect loss of catalytic or substrate-recognition function rather than protein instability. Together, these results identify K118, D184, and W188 as key residues that shape the Dag1 active site and suggest that it employs a canonical glycosylase mechanism in which a lysine–aspartate catalytic pair and a base-stacking tryptophan coordinate substrate cleavage.

### Phage escape from Dag1 confirms evolutionary trade-off

To determine whether phages could evolve resistance to Dag1, we focused on HK243, which encodes a well-defined gene cluster for dPreQ_0_ biosynthesis and incorporation (Fig. 1E). When HK243 was plated on *E. coli* expressing Dag1, rare plaques appeared after overnight incubation, indicating successful infection despite the presence of the defence system. Individual plaques were isolated and re-tested on Dag1-expressing cells, confirming that these mutants bypassed defence (Fig. 2E). Whole-genome sequencing of three escape mutants revealed an identical point mutation in *dpdA*, resulting in a G217E substitution in the transglycosylase protein that incorporates dPreQ_0_ into DNA. AlphaFold3 modelling suggested that this substitution disrupts the positively charged DNA-binding surface of DpdA (*26*) (Fig. 2F and G). HPLC-MS analysis of the escaper phage genome confirmed complete loss of dPreQ_0_ incorporation (Fig. S2A and Table 1), and an *in vitro* cleavage assay showed that Dag1 no longer degraded its DNA (Fig. 2A).

We next tested whether the loss of dPreQ_0_ modification in the Dag1 escaper phage altered its susceptibility to host restriction enzymes. Genomic DNA from wild type HK243 resisted cleavage by enzymes with GC-containing recognition sites (EcoRV, HaeIII, NdeI), whereas it was efficiently digested by SspI, which has an AT-rich recognition site (Fig. 2H). By contrast, DNA from the HK243 escape mutant was efficiently cleaved by all four enzymes under the same conditions (Fig. 2H). These findings confirm that guanine modifications protect HK243 DNA from host restriction enzymes with CG-containing recognition sites and that escaper mutants regain sensitivity once the modification is lost. Together, these results demonstrate that HK243 can evade Dag1 activity by abandoning dPreQ_0_ modification, but this escape comes at the cost of susceptibility to host restriction enzymes. This trade-off highlights the evolutionary tension between restriction evasion through genome modification and glycosylase defences that exploit these same chemical signatures.

### Structure-guided discovery reveals diverse integron-encoded glycosylases

Dag1 could not be linked to known DNA glycosylases by sequence similarity, yet it clearly provided defence against phages with modified DNA. This raised the possibility that additional defence glycosylases exist that similarly escape sequence-based detection. Identifying such proteins across entire bacterial genomes is challenging because every species encodes numerous housekeeping glycosylases that share conserved catalytic motifs but differ in function, making defence-associated variants difficult to distinguish by homology alone. To overcome this limitation, we focused our search within sedentary chromosomal integrons in Vibrionaceae, which are increasingly recognized as major reservoirs of antiviral defence systems (*21*, *27*, *28*). These elements are uniquely suited for targeted discovery: they consist entirely of horizontally transferred DNA, their gene cassettes are structurally delimited by conserved recombination sites, and they are highly enriched for defence genes relative to the rest of the genome. In *V. parahaemolyticus*, integrons account for only 1–2% of the genome yet encode ∼11% of identifiable defence systems, and a recent screen showed that ∼20% of cassettes provide phage resistance (*21*). Together, these features make integrons an ideal and tractable platform for uncovering streamlined antiviral effectors invisible to conventional sequence-based homology searches.

To systematically search for additional glycosylase-like proteins, we assembled a database of predicted protein structures from complete integrons identified in 1,094 non-redundant Vibrionaceae genomes. This dataset comprised 907 integrons encoding 76,370 proteins, which we clustered into 8,999 families based on sharing ≥30% pairwise identity across ≥80% of the sequence. Structures were predicted for representative members of each family using ESM3 (*29*), and the resulting structures were queried against Dag1 and Dag2 predicted structural models using FoldSeek (*30*). This search identified 24 structural clusters with high-confidence similarity to Dag1 or Dag2, including their own families and 22 new groups. Together, these clusters encompassed 109 proteins that shared a common fold despite low sequence identity, typically below 20% (fig. S4A and B). Structural alignments using DALI (*31*) revealed a common catalytic architecture and predicted DNA-binding pocket across candidates, even among the most sequence-diverse proteins (Fig. 3A, fig. S4C).

**Fig. 3.**
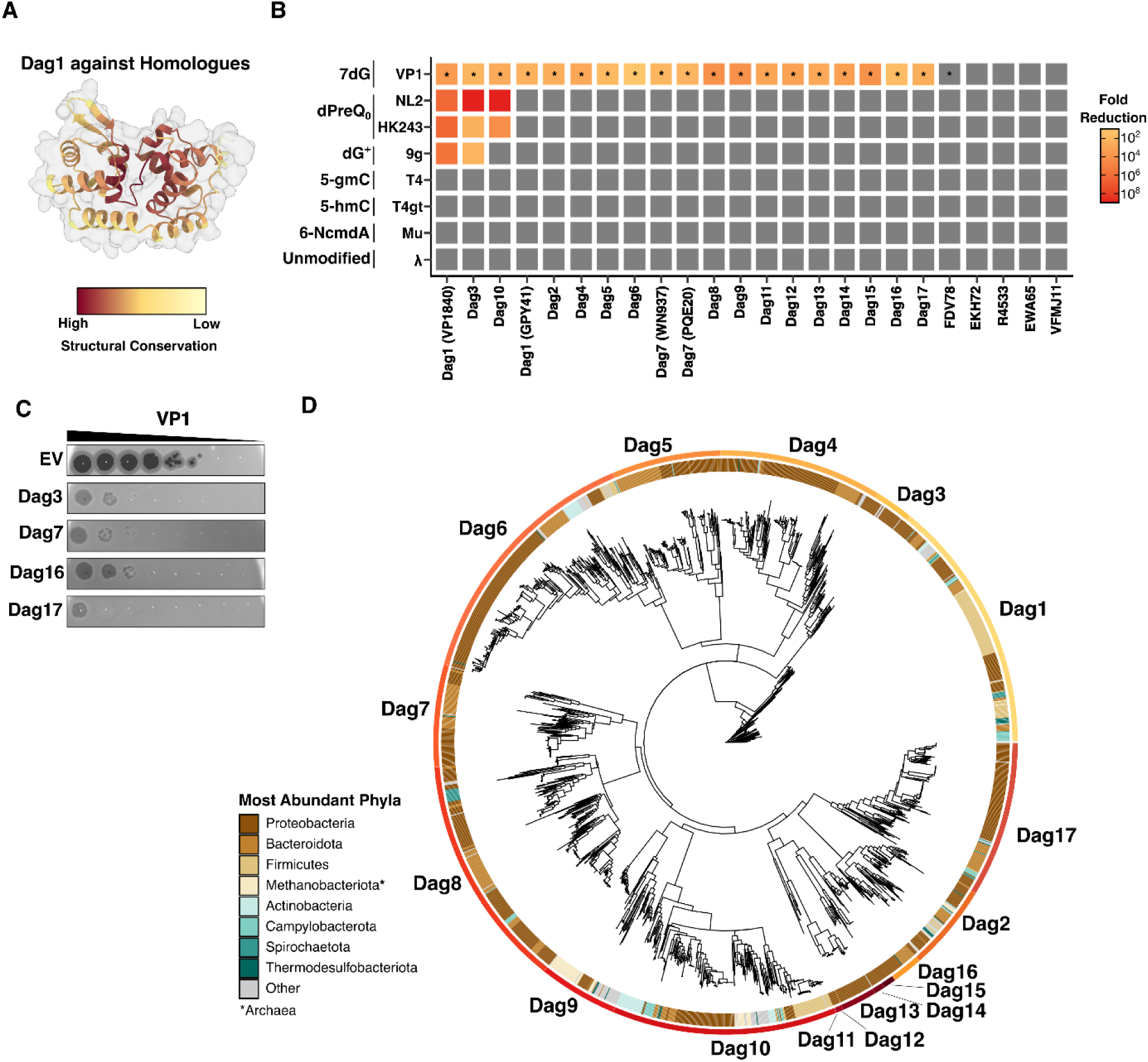
Integron-encoded Dag structural analogs also provide phage defence. **(A)** All 23 additional homologues were aligned by structure to Dag1 using ChimeraX, average RMSD values for each position were calculated, and the cartoon structure was coloured by this value. Dark red indicates low RMSD values and yellow indicates high RMSD values for a given position. (B) Phage plating assays of each homologue tested were tested against VP1 (7dG-modified), NL2 and HK243 (dPreQo), 9g (dG+), T4 (5-gmc) and T4gt (5-hmC), Mu (6-NcmdA), and λ and compared to empty vector controls to generate a fold-reduction matrix. Fold-reduction was calculated from biological duplicates. Matrix is represented as a heatmap. Asterisks indicate reduced plaque size or otherwise modified plaque morphology. (C) Representative phage plating assays of select Dag family members against VP 1 shown in **(B). (D)** Maximum-likelihood phylogenetic tree of Dag homologs as identified by PSI-BLAST filtered by sequence identity (>30%) and coverage (>70%). Only Dag homologues which provided antiphage function are shown and individual Dag clades are shown and are labelled. Phlya are indicated for each tree tip.

To test whether these proteins with predicted glycosylase folds mediate antiviral activity, we selected one representative from each of the 22 newly identified clusters (table S3) and assessed their ability to protect *V. parahaemolyticus* and *E. coli* against a panel of phages with distinct DNA modifications. Expression of 17 candidates markedly reduced the plating efficiency of the 7-deazaguanine modified Vibriophage VP1 (Fig. 3B). However, like Dag2, most failed to protect against the preQ_0_ modified phage NL2 or the *E. coli* phages HK243 and 9g, which contain preQ_0_ and G modifications, respectively. None provided defence against phages carrying unrelated DNA modifications, like T4, T4*gt*, or Mu, nor against unmodified phage λ (Fig. 3B and C).

We next examined the evolutionary distribution of sequence homologues of the glycosylases with confirmed antiphage activity. Iterative PSI-BLAST searches using the protein sequences of the 19 validated defence glycosylases identified homologues across thousands of bacterial genomes spanning diverse phyla, including Proteobacteria, Bacteroidota, Firmicutes, Actinobacteria, and Campylobacterota, among others (Fig. 3D, table S4 and S5). Pairwise similarity network analysis (fig. S5) revealed that while certain families shared overlapping homologues, for example, Dag1 (VP1840 and HF305) and Dag7 (PQE20 and WN937), most formed distinct, cohesive clusters, consistent with their diversification into separate functional lineages. Taken together, these results show that *Vibrio* integrons harbour a remarkably diverse and abundant repertoire of glycosylases that selectively target 7-deazaguanine modified phages, revealing an extensive layer of glycosylase-driven antiviral immunity.

### From genome maintenance to immunity: the repurposing of glycosylase folds for defence

To place the defence-associated glycosylases in evolutionary context, we compared them with canonical bacterial DNA glycosylases involved in base excision repair. Across 80 complete *Vibrio* reference genomes, we identified six highly conserved DNA glycosylases (Fig. 4A) that are well-established components of bacterial base excision repair pathways (*19*). Members of each family shared high sequence identity and were almost universally present in the *Vibrio* core genome (table S6), reflecting their stable integration and conservation across species. By contrast, Dag-like glycosylases were frequently encoded within integrons or adjacent to mobile genetic elements. On average, ∼45% of Dag homologues were located within 20 kB of a known defence-associated gene or mobile element marker such as a transposase, recombinase, or integrase, compared with only 3.1% of the housekeeping glycosylases (table S7 and 8). This strong association with mobile elements suggests that defence-associated glycosylases are actively exchanged among bacterial populations through horizontal gene transfer. Because horizontally acquired genes often display compositional sequence biases, we also measured the local G/C content surrounding each glycosylase. Dags were enriched in atypical, A/T-rich regions of the genome (average ΔG/C = −2.37%), whereas housekeeping glycosylases showed no such deviation (Fig. 4B and table S8). Together, these observations indicate that glycosylases co-opted for defence occupy a dynamic and mobilizable evolutionary niche distinct from their chromosomally anchored DNA repair counterparts.

**Fig 4.**
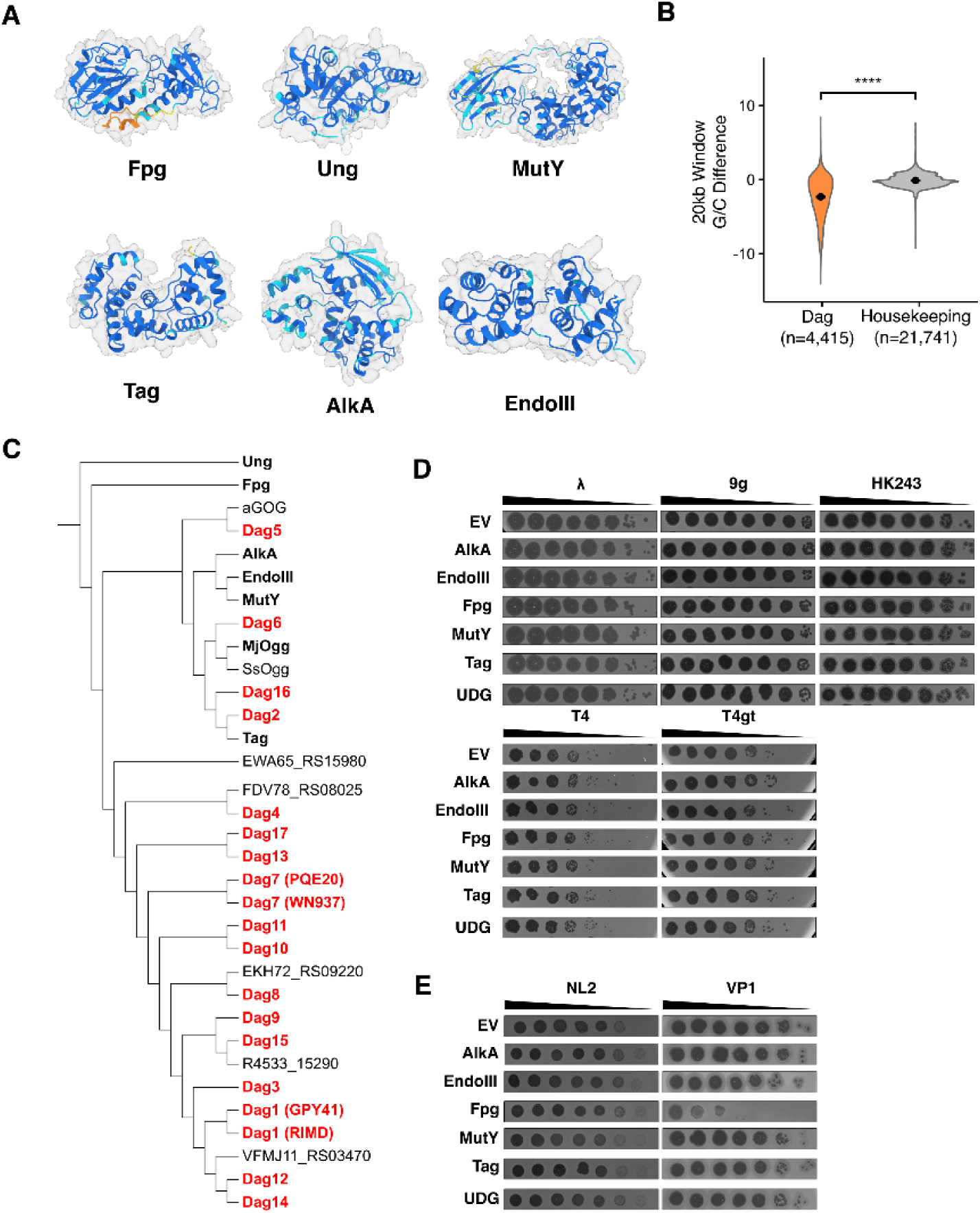
Dag families arc distinct from housekeeping glyeosylascs by genomic context, protein structure, and antiphage function. **(A)** AlphaFold3 predicted structures of RTMD 2210633 encoded housekeeping glycosylases. Full-length proteins are shown, with the exception of AlkA, in which only the glycosylase domain is shown (AA sequence 200-454). AlphaFold3 structures are coloured by pLDDT at each residue (dark blue, pLDDT >90; light blue, 90 > pLDDT > 70; yellow, low 70 > pLDDT > 50; orange, very low pLDDT<50). **(B)** ¾GC was calculated for both 20kb windows surrounding either housekeeping or Dag glycosylases and compared to genome G/C content. G/C content differences are plotted and n values represent number of genome windows analyzed. A welch’s t-test was performed, **** = p<0.0005. **(C)** DALI structural alignments were performed in all-against-all mode, and the resulting structural tree is shown. Housekeeping glycosylases shown in **(A)** are balded, and Dag glycosylases with described antiphage function are indicated in red. Structures for aGOG (7OLB), ssOgg (3FHG), and mjOgg (3KNT) were obtained from the PDB, the remainder were AlphaFold3 predicted structures. **(D)** Phage plating assay of expressed housekeeping glycosylases against coliphages A, 9g, HK243, T4, and *T4gt.* **(E)** Same as in **(D),** except with Vibriophages NL2 (RTMD 2210633) and VPl (VP3283-61).

To explore whether these differences extend to protein structure and evolutionary history, we compared defence-associated and housekeeping glycosylases using both sequence-based and structure-guided phylogenetic analyses. Sequence clustering showed that bacterial housekeeping and defence-associated glycosylases form distinct, non-overlapping families across *Vibrio* species (fig. S5). As the extremely high sequence variation we observed obscured deeper evolutionary relationships among families, we next used fold-space clustering to assess protein structural relatedness among defence-associated and housekeeping glycosylases.

Structurally, bacterial DNA glycosylases fall into four major superfamilies defined by their catalytic folds (table S1). DALI-based structural analysis revealed that all defence-active Dag proteins belong to the HhH superfamily, but they have a polyphyletic distribution within it rather than forming a single lineage (Fig. 4C). Most Dags cluster together in a branch distinct from housekeeping enzymes, indicating that they have diverged substantially from the canonical repair enzymes. However, several defence-active enzymes are interspersed among bacterial repair glycosylases, suggesting that glycosylase-based defence has arisen multiple times through independent co-option of repair proteins for antiviral function.

The clustering of some Dags near canonical bacterial repair enzymes like Tag and aGOG (Fig. 4C) raised the question whether housekeeping glycosylases with similar folds can also mediate phage defence. To address this, we cloned one representative from each of the six core families and tested their ability to inhibit phage replication. None of these enzymes provided measurable protection against phages in *E. coli* (Fig. 4D). In *V. parahaemolyticus*, however, overexpression of Fpg conferred ∼100-fold protection against phage VP1, which is fully modified with 7-deazagunaine (Fig. 4E). It had no effect on NL2, which carries dPreQ_0_ and is only partially modified. Fpg normally removes oxidized purines like 8-oxoguanine from host DNA (*32*, *33*), so its weak antiviral activity may reflect partial recognition of chemically similar guanine derivatives in modified phage genomes. This expression-dependent and phage-specific protection suggests a latent ability of Fpg to recognize certain noncanonical bases, supporting the idea that basal repair activities can be co-opted and refined for antiviral defence. Overall, these results emphasize the separation between housekeeping and defence-associated glycosylases.

Whereas canonical enzymes are optimized for genome maintenance, Dag proteins represent a distinct, mobilizable class of antiviral effectors. Their broad distribution and consistent enrichment within defence loci highlight how glycosylases have been repeatedly adapted as a widespread molecular strategy for recognizing and destroying chemically modified phage DNA.

### Integrons also encode UDG-like proteins with antiphage activity

The discovery of Brig1 (*18*), a UDG-family enzyme that functions in antiviral defence by excising modified cytosines from the genomes of T-even phages, shows that glycosylases outside the HhH superfamily can also be repurposed for defence. Building on this observation, we next asked whether integrons also encode divergent UDG or Helix Two-Turn Helix (H2TH) family enzymes (table S1). This analysis recovered 27 protein sequences with high-confidence structural similarity to uracil DNA glycosylases. These candidates, each sharing ≤20% sequence identity with Brig1, grouped into 11 distinct sequence clusters at a 30% identity cutoff (fig. S6A, S6B, and table S9). The most consistent feature across these UDG-like proteins was a conserved β-sheet core flanked by helices characteristic of uracil DNA glycosylases (Fig. 5A and B). In contrast to Dags, however, the predicted DNA-binding pocket was not strongly positionally conserved (Fig. 3A and 5A).

**Fig 5.**
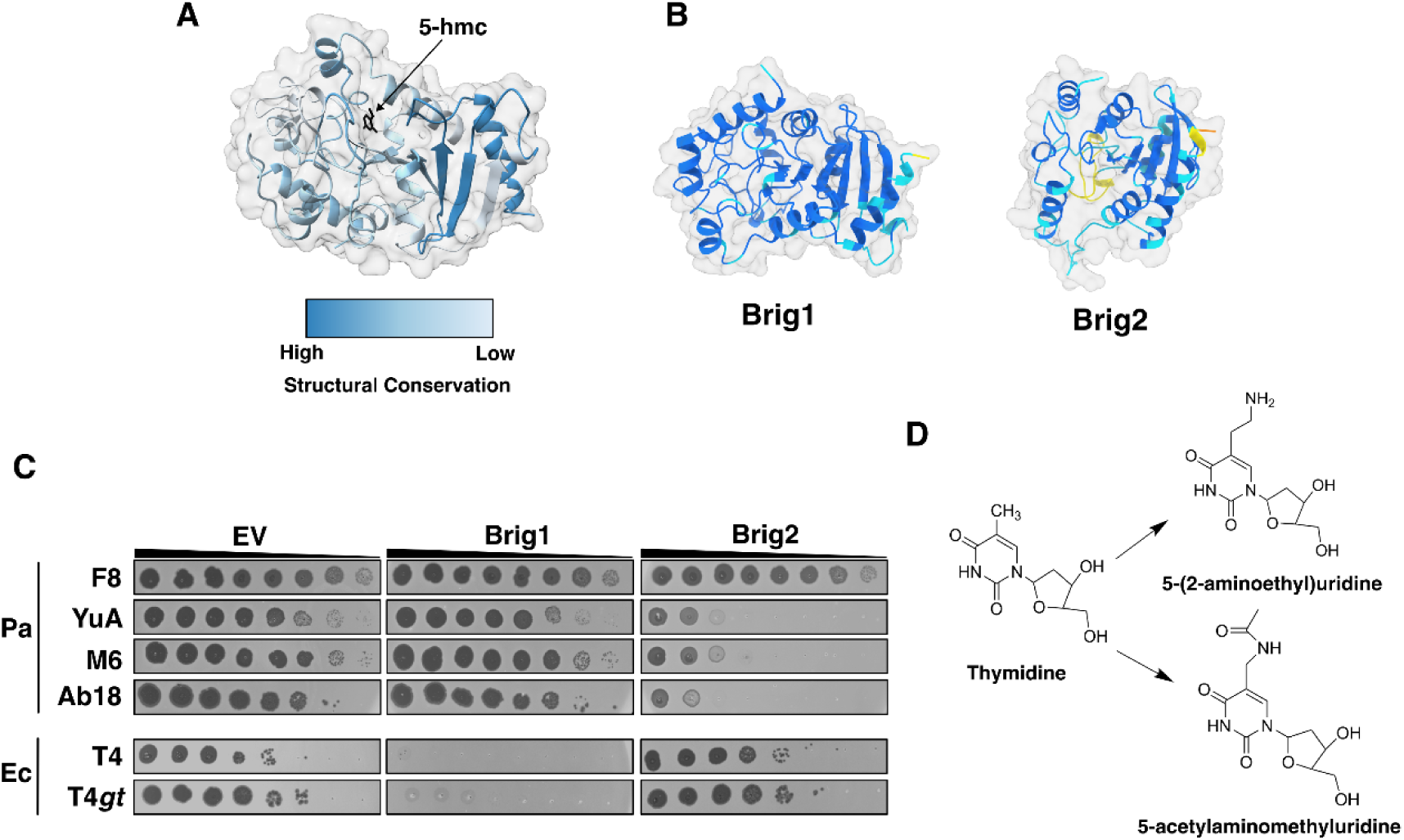
A UDG structural homologue provides defence against thymidine modified phages. **(A)** Brigl structural homologues were aligned to the Brigl structure using ChimeraX and average RMSD values were calculated for each residue position. The Brigl cartoon structure is shown and coloured by average RMSD values at each position. 5-hydroxymethylcytosine is modelled into its binding pocket and coloured in black. **(B)** AlphaFold3 predicted structures of Brigl and Brig2 are shown, coloured by pLDDT by AlphaFold3 colouring. Foldseek E-value and RMSD for the alignment of these two proteins was 2.453E-5 and 7.342 respectively. **(C)** Phage plating experiments for empty vector (EV) and cells expressing either Brigl or Brig2 are shown. Experiments were done in both *P. aeruginosa* (Pa) and in *E. coli* (Ec) as labelled. Images are representative of three biological replicates. (D) Schematic diagram of phage M6/YuA thymidine modification to 5-(2-aminoethyl)uridine and phage Ab 18 thymidine modification to 5-acetylaminomethy 1-2 ’-deoxyuridine.

To test whether these UDG proteins mediate antiphage defence, we selected seven candidates (table S10) and assayed their ability to inhibit phage replication in *E. coli*, *V. parahaemolyticus*, and *P. aeruginosa*. None provided defence against T4 or other phages encoding modified cytosine residues like Brig1 does, or against phages with guanine modifications targeted by the Dags. However, one candidate conferred strong defence against *Pseudomonas* phages M6 and YuA, which incorporate 5-(2-aminoethyl)uridine into their genomes (*34–36*), as well as phage Ab18, which carries a different thymidine derivative, 5-acetylaminomethyl-2′-deoxyuridine (*36*) (Fig. 5C and D). We designated this enzyme Brig2.

A PSI-BLAST search using Brig2 as a query identified more than 500 homologues across diverse bacterial phyla, including Proteobacteria, Bacteroidetes and Firmicutes (tables S4 and S5). On average, 41% of Brig2-like proteins were encoded within 20 kb of a mobile genetic element such as a transposase, recombinase, or integrase, or near known defence systems (table S7 and 8). This widespread association with mobile elements suggests that these UDG-family enzymes are actively disseminated through horizontal gene transfer. Genome composition analysis further revealed that Brig2 homologues, like Dag family members, are preferentially situated in A/T-rich regions that bear hallmarks of recent acquisition and defence island enrichment (average ΔG/C = −1.70; tables S7 and 8). Together, these findings indicate that UDG proteins have been independently co-opted for phage defence. More broadly, the discovery of both guanine- and thymidine-targeting defence glycosylases underscores the value of structure-based discovery for revealing mechanistically related antiviral enzymes that escape sequence-based detection.

## Discussion

Our findings establish that DNA glycosylases, best known for their roles in DNA repair, also constitute a widespread and diverse class of antiviral effectors. Using structure-guided discovery, we identified 18 integron-encoded glycosylase families that mediate defence against phages carrying chemically modified bases, including guanine and thymidine derivatives. These defence-associated enzymes differ from housekeeping DNA repair glycosylases in that they are encoded within mobile genomic regions and show sporadic phylogenetic distributions consistent with horizontal transfer. The glycosylase fold is inherently suited to this defensive role, having evolved to recognize and excise a broad range of chemically altered bases. This versatility likely contributes to its repeated recruitment for defence against phages with modified DNA.

The widespread occurrence of defence-associated glycosylases, together with type IV restriction enzymes that recognize modified bases absent from host DNA, demonstrate that modification-dependent recognition has repeatedly evolved as a strategy for distinguishing self from non-self during infection. While type IV restriction enzymes separate DNA binding and cleavage into distinct domains on the same protein, or even into different proteins, glycosylases function as compact single-domain enzymes that both recognize and excise the modified base. This compact architecture is particularly well suited to *Vibrio* integrons, where gene cassettes are constrained in size and multi-component systems are less easily accommodated. As the characterized diversity of phage genome modifications continues to expand, additional enzymatic strategies co-opted to exploit these chemical signatures are likely to be uncovered.

The emergence of antiviral glycosylases from housekeeping repair enzymes illustrates how ancient enzymatic folds can be repeatedly co-opted for immunity. The weak, phage-specific protection conferred by overexpressed Fpg suggests that recognition of chemically similar modified bases can provide a foothold for the evolution of antiviral glycosylases. Comparative analyses support this view, showing that defence-associated glycosylases occupy multiple positions within the HhH structural superfamily rather than a single lineage. Several active defence enzymes are interspersed among canonical repair proteins, implying that glycosylase-based defence has emerged independently multiple times through separate co-option events.

Brig1 and Brig2, which belong to the uracil DNA glycosylase family but act on cytosine- and thymidine-modified phages, represent an independent occurrence of the same defence principle in a distinct fold class. Viewed together, these findings demonstrate that proteins with glycosylase-like folds have evolved antiviral functions on several occasions, representing a recurring solution to the problem of detecting and degrading chemically disguised phage DNA.

Across evolution, DNA base modifications have emerged as focal points of host-virus conflict with glycosylases mobilized in diverse contexts. In bacteria, glycosylases transform phage DNA modifications into defence triggers. In eukaryotic viruses, DNA glycosylases play opposing roles in virus-host conflict. Herpesviruses encode UNG glycosylases to maintain genome integrity during both active replication and latency (*37*), while in HIV-1, deamination of viral DNA by the host APOBEC3G cytidine deaminase provides an innate barrier to infection (*38–42*). To counter this, the virus packages the host UNG2 glycosylase into virions to remove uracils during reverse transcription and encodes the Vif protein to exclude the cytidine deaminase APOBEC3G from viral particles (*43*, *44*). A comparable interplay may also occur in bacteria, where defence-associated glycosylases detect phage DNA modifications as antiviral triggers, and phages could evolve countermeasures to evade or repair glycosylase-mediated damage. Such countermeasures could include inhibitors that are pre-packaged within the virion and injected alongside the genome, analogous to the IPI* protein of phage T4 that blocks modification-dependent restriction systems (*45*). Other strategies could involve altering the chemistry of modified bases, encoding protective DNA-binding proteins, or recruiting repair enzymes to reverse excision events. Together, these examples show that DNA glycosylases have become key participants in virus-host conflicts across domains of life, sometimes mobilized in defence and other times co-opted by viruses for protection.

Finally, this work highlights the power of structure-guided discovery to uncover cryptic components of bacterial immunity. By revealing distant evolutionary relationships, structural approaches can expose an uncharted layer of microbial immunity built on enzymatic scaffolds that can be repurposed to counter viral innovation. As predictive accuracy improves and structural databases continue to expand, this strategy will enable systematic mapping of microbial immunity and illuminate how ancient enzyme folds are reinvented to meet viral challenge.

## Acknowledgements

The authors thank Dr. Alan Davidson, Dr. Haley Wyatt, and members of the Maxwell Lab for thoughtful discussion and comments on this article. The authors also thank Dr. Joseph Bondy-Denomy for providing the *Psuedomonas* phages F8, YuA, M6, and Ab18, and Pramalkumar Patel for sharing phage PHP2.

## Funding

This study was supported by grants from the Canadian Institutes of Health Research (PJT-165936), the Natural Sciences and Engineering Research Council (RGPIN-2023-05366; SMFSU-581368-2023), and the Canadian Foundation for Innovation/Ontario Research Fund (44010) to K.L.M. L.J.G. is supported by a Postdoctoral Fellowship from the Canadian Institutes of Health Research and the GSK EPIC Convergence Postdoctoral Fellowship in Antimicrobial Resistance granted by the Emerging Pandemic and Infections Consortium (EPIC) at the University of Toronto. K.L.M. is the Canada Research Chair in Bacteriophage Biology and Therapeutics (CRC-2023-00010).

### Author contributions

Conceptualization: LJG, ALQ, PRW, KLM

Methodology: LJG, ALQ, MSB, Y-JL, PRW, KLM

Investigation: LJG, ALQ, YVL, SRF, MSB, Y-JL, PRW, KLM

Visualization: LJG, ALQ, Y-JL, KLM

Funding acquisition: LJG, KLM

Project administration: LJG, KLM

Supervision: LJG, PRW, KLM

Writing – original draft: LJG, ALQ, KLM

Writing – review & editing: LJG, ALQ, YVL, SRF, MSB, Y-JL, PRW, KLM

## Data and materials availability

All phage, plasmid, and strain information is available in tables S10 and S11. Genome sequences for phage VP1, VP12, HK243, NL2, and PHP2 have been deposited to the NCBI genome database and will be available upon publication of the manuscript.

## Supplementary Materials

Materials and Methods

Figs. S1 to S6

Tables S1 to S11

References (46–58)

## References and Notes

1. W. A. M. Loenen, D. T. F. Dryden, E. A. Raleigh, G. G. Wilson, N. E. Murray, Highlights of the DNA cutters: a short history of the restriction enzymes. Nucleic Acids Res 42, 3–19 (2014).

2. H. Georjon, A. Bernheim, The highly diverse antiphage defence systems of bacteria. Nat Rev Microbiol 21, 686–700 (2023).

3. K. S. Makarova, Y. I. Wolf, J. Iranzo, S. A. Shmakov, O. S. Alkhnbashi, S. J. J. Brouns, E. Charpentier, D. Cheng, D. H. Haft, P. Horvath, S. Moineau, F. J. M. Mojica, D. Scott, S. A. Shah, V. Siksnys, M. P. Terns, Č. Venclovas, M. F. White, A. F. Yakunin, W. Yan, F. Zhang, R. A. Garrett, R. Backofen, J. van der Oost, R. Barrangou, E. V. Koonin, Evolutionary classification of CRISPR–Cas systems: a burst of class 2 and derived variants. Nat Rev Microbiol 18, 67–83 (2020).

4. C. Wang, Z. Shen, X.-Y. Yang, T.-M. Fu, Structures and functions of short argonautes. RNA Biology 21, 883–889 (2024).

5. C. N. Vassallo, C. R. Doering, M. L. Littlehale, G. I. C. Teodoro, M. T. Laub, A functional selection reveals previously undetected anti-phage defence systems in the *E. coli* pangenome. Nat Microbiol 7, 1568–1579 (2022).

6. L. Rodriguez-Rodriguez, J. Pfister, L. Schuck, A. E. Martin, L. M. Mercado-Santiago, V. S. Tagliabracci, K. J. Forsberg, Metagenomic selections reveal diverse antiphage defenses in human and environmental microbiomes. Cell Host & Microbe 33, 1381–1395.e7 (2025).

7. Y. Cui, Z. Dai, Y. Ouyang, C. Fu, Y. Wang, X. Chen, K. Yang, S. Zheng, W. Wang, P. Tao, Z. Guan, T. Zou, Bacterial Hachiman complex executes DNA cleavage for antiphage defense. Nat Commun 16, 2604 (2025).

8. O. T. Tuck, B. A. Adler, E. G. Armbruster, A. Lahiri, J. J. Hu, J. Zhou, J. Pogliano, J. A. Doudna, Genome integrity sensing by the broad-spectrum Hachiman antiphage defense complex. Cell 187, 6914–6928.e20 (2024).

9. Y. Gu, H. Li, A. Deep, E. Enustun, D. Zhang, K. D. Corbett, Bacterial Shedu immune nucleases share a common enzymatic core regulated by diverse sensor domains. Molecular Cell 85, 523–536.e6 (2025).

10. G. Hutinet, W. Kot, L. Cui, R. Hillebrand, S. Balamkundu, S. Gnanakalai, R. Neelakandan, A. B. Carstens, C. Fa Lui, D. Tremblay, D. Jacobs-Sera, M. Sassanfar, Y.-J. Lee, P. Weigele, S. Moineau, G. F. Hatfull, P. C. Dedon, L. H. Hansen, V. de Crécy-Lagard, 7-Deazaguanine modifications protect phage DNA from host restriction systems. Nat Commun 10, 5442 (2019).

11. L. Cui, S. Balamkundu, C.-F. Liu, H. Ye, J. Hourihan, A. Rausch, C. Hauß, E. Nilsson, M. Hoetzinger, K. Holmfeldt, W. Zhang, L. Martinez-Alvarez, X. Peng, D. Tremblay, S. Moinau, N. Solonenko, M. B. Sullivan, Y.-J. Lee, A. Mulholland, P. R. Weigele, V. de Crécy-Lagard, P. C. Dedon, G. Hutinet, Four additional natural 7-deazaguanine derivatives in phages and how to make them. Nucleic Acids Res 51, 9214–9226 (2023).

12. R. L. Sinsheimer, Nucleotides from T2r+ Bacteriophage. Science 120, 551–553 (1954).

13. I. R. Lehman, E. A. Pratt, On the Structure of the Glucosylated Hydroxymethylcytosine Nucleotides of Coliphages T2, T4, and T6. Journal of Biological Chemistry 235, 3254–3259 (1960).

14. G. Hutinet, Y.-J. Lee, V. de Crécy-Lagard, P. R. Weigele, Hypermodified DNA in Viruses of *E. coli* and *Salmonella*. EcoSal Plus 9, eESP-0028-2019 (2021).

15. E. A. Raleigh, G. Wilson, *Escherichia coli* K-12 restricts DNA containing 5-methylcytosine. Proc Natl Acad Sci U S A 83, 9070–9074 (1986).

16. T. A. Bickle, D. H. Krüger, Biology of DNA restriction. Microbiol Rev 57, 434–450 (1993).

17. C. L. Bair, L. W. Black, A Type IV modification dependent restriction nuclease that targets glucosylated hydroxymethyl cytosine modified DNAs. J Mol Biol 366, 768–778 (2007).

18. A. A. Hossain, Y. Z. Pigli, C. F. Baca, S. Heissel, A. Thomas, V. K. Libis, J. Burian, J. S. Chappie, S. F. Brady, P. A. Rice, L. A. Marraffini, DNA glycosylases provide antiviral defence in prokaryotes. Nature 629, 410–416 (2024).

19. A. L. Jacobs, P. Schär, DNA glycosylases: in DNA repair and beyond. Chromosoma 121, 1–20 (2012).

20. C. H. Trasviña-Arenas, M. Demir, W.-J. Lin, S. S. David, Structure, function and evolution of the Helix-hairpin-Helix DNA glycosylase superfamily: Piecing together the evolutionary puzzle of DNA base damage repair mechanisms. DNA Repair (Amst) 108, 103231 (2021).

21. L. J. Getz, S. R. Fairburn, Y. Vivian Liu, A. L. Qian, K. L. Maxwell, Integrons are antiphage defence libraries in *Vibrio parahaemolyticus*. Nat Microbiol 10, 724–733 (2025).

22. J. J. Thiaville, S. M. Kellner, Y. Yuan, G. Hutinet, P. C. Thiaville, W. Jumpathong, S. Mohapatra, C. Brochier-Armanet, A. V. Letarov, R. Hillebrand, C. K. Malik, C. J. Rizzo, P. C. Dedon, V. de Crécy-Lagard, Novel genomic island modifies DNA with 7-deazaguanine derivatives. Proceedings of the National Academy of Sciences 113, E1452–E1459 (2016).

23. C. S. Crippen, Y.-J. Lee, G. Hutinet, A. Shajahan, J. C. Sacher, P. Azadi, V. de Crécy-Lagard, P. R. Weigele, C. M. Szymanski, Deoxyinosine and 7-Deaza-2-Deoxyguanosine as Carriers of Genetic Information in the DNA of *Campylobacter* Viruses. J Virol 93, e01111–19 (2019).

24. S. R. Johnson, P. R. Weigele, A. Fomenkov, A. Ge, A. Vincze, J. B. Eaglesham, R. J. Roberts, Z. Sun, Domainator, a flexible software suite for domain-based annotation and neighborhood analysis, identifies proteins involved in antiviral systems. Nucleic Acids Res 53, gkae1175 (2025).

25. A. Sarre, M. Ökvist, T. Klar, D. R. Hall, A. O. Smalås, S. McSweeney, J. Timmins, E. Moe, Structural and functional characterization of two unusual endonuclease III enzymes from *Deinococcus radiodurans*. Journal of Structural Biology 191, 87–99 (2015).

26. S. H. Gedara, E. Wood, A. Gustafson, C. Liang, S.-H. Hung, J. Savage, P. Phan, A. Luthra, V. de Crécy-Lagard, P. Dedon, M. A. Swairjo, D. Iwata-Reuyl, 7-Deazaguanines in DNA: functional and structural elucidation of a DNA modification system. Nucleic Acids Res 51, 3836–3854 (2023).

27. B. Darracq, E. Littner, M. Brunie, J. Bos, P. A. Kaminski, F. Depardieu, W. Slesak, K. Debatisse, M. Touchon, A. Bernheim, D. Bikard, F. Le Roux, D. Mazel, E. P. C. Rocha, C. Loot, Sedentary chromosomal integrons as biobanks of bacterial antiphage defense systems. Science 388, eads0768 (2025).

28. N. Kieffer, A. Hipólito, L. Ortiz-Miravalles, P. Blanco, T. Delobelle, P. Vizuete, F. M. Ojeda, T. Jové, D. Jurenas, M. García-Quintanilla, A. Carvalho, P. Domingo-Calap, J. A. Escudero, Mobile integrons encode phage defense systems. Science 388, eads0915 (2025).

29. T. Hayes, R. Rao, H. Akin, N. J. Sofroniew, D. Oktay, Z. Lin, R. Verkuil, V. Q. Tran, J. Deaton, M. Wiggert, R. Badkundri, I. Shafkat, J. Gong, A. Derry, R. S. Molina, N. Thomas, Y. A. Khan, C. Mishra, C. Kim, L. J. Bartie, M. Nemeth, P. D. Hsu, T. Sercu, S. Candido, A. Rives, Simulating 500 million years of evolution with a language model. Science 387, 850–858 (2025).

30. M. van Kempen, S. S. Kim, C. Tumescheit, M. Mirdita, J. Lee, C. L. M. Gilchrist, J. Söding, M. Steinegger, Fast and accurate protein structure search with Foldseek. Nat Biotechnol 42, 243–246 (2024).

31. L. Holm, Dali server: structural unification of protein families. Nucleic Acids Res 50, W210–W215 (2022).

32. S. Boiteux, T. R. O’Connor, J. Laval, Formamidopyrimidine DNA glycosylase of *Escherichia coli*: cloning and sequencing of the *fpg* structural gene and overproduction of the protein. The EMBO Journal 6, 3177–3183 (1987).

33. J. Tchou, V. Bodepudi, S. Shibutani, I. Antoshechkin, J. Miller, A. P. Grollman, F. Johnson, Substrate specificity of Fpg protein. Recognition and cleavage of oxidatively damaged DNA. Journal of Biological Chemistry 269, 15318–15324 (1994).

34. P.-J. Ceyssens, V. Mesyanzhinov, N. Sykilinda, Y. Briers, B. Roucourt, R. Lavigne, J. Robben, A. Domashin, K. Miroshnikov, G. Volckaert, K. Hertveldt, The Genome and Structural Proteome of YuA, a New *Pseudomonas aeruginosa* Phage Resembling M6. Journal of Bacteriology 190, 1429–1435 (2008).

35. Y.-J. Lee, N. Dai, S. E. Walsh, S. Müller, M. E. Fraser, K. M. Kauffman, C. Guan, I. R. Corrêa, P. R. Weigele, Identification and biosynthesis of thymidine hypermodifications in the genomic DNA of widespread bacterial viruses. Proceedings of the National Academy of Sciences 115, E3116–E3125 (2018).

36. Y.-J. Lee, N. Dai, S. I. Müller, C. Guan, M. J. Parker, M. E. Fraser, S. E. Walsh, J. Sridar, A. Mulholland, K. Nayak, Z. Sun, Y.-C. Lin, D. G. Comb, K. Marks, R. Gonzalez, D. P. Dowling, V. Bandarian, L. Saleh, I. R. Corrêa, P. R. Weigele, Pathways of thymidine hypermodification. Nucleic Acids Res 50, 3001–3017 (2022).

37. J. Sire, G. Quérat, C. Esnault, S. Priet, Uracil within DNA: an actor of antiviral immunity. Retrovirology 5, 45 (2008).

38. R. S. Harris, K. N. Bishop, A. M. Sheehy, H. M. Craig, S. K. Petersen-Mahrt, I. N. Watt, M. S. Neuberger, M. H. Malim, DNA deamination mediates innate immunity to retroviral infection. Cell 113, 803–809 (2003).

39. D. Lecossier, F. Bouchonnet, F. Clavel, A. J. Hance, Hypermutation of HIV-1 DNA in the absence of the Vif protein. Science 300, 1112 (2003).

40. R. Mariani, D. Chen, B. Schröfelbauer, F. Navarro, R. König, B. Bollman, C. Münk, H. Nymark-McMahon, N. R. Landau, Species-specific exclusion of APOBEC3G from HIV-1 virions by Vif. Cell 114, 21–31 (2003).

41. A. F. Weil, D. Ghosh, Y. Zhou, L. Seiple, M. A. McMahon, A. M. Spivak, R. F. Siliciano, J. T. Stivers, Uracil DNA glycosylase initiates degradation of HIV-1 cDNA containing misincorporated dUTP and prevents viral integration. Proc Natl Acad Sci U S A 110, E448–E457 (2013).

42. B. Yang, K. Chen, C. Zhang, S. Huang, H. Zhang, Virion-associated uracil DNA glycosy-lase-2 and apurinic/apyrimidinic endonuclease are involved in the degradation of APO-BEC3G-edited nascent HIV-1 DNA. J Biol Chem 282, 11667–11675 (2007).

43. S. Priet, N. Gros, J.-M. Navarro, J. Boretto, B. Canard, G. Quérat, J. Sire, HIV-1-associated uracil DNA glycosylase activity controls dUTP misincorporation in viral DNA and is essential to the HIV-1 life cycle. Mol Cell 17, 479–490 (2005).

44. S. Priet, J.-M. Navarro, N. Gros, G. Quérat, J. Sire, Differential incorporation of uracil DNA glycosylase UNG2 into HIV-1, HIV-2, and SIV(MAC) viral particles. Virology 307, 283–289 (2003).

45. D. Rifat, N. T. Wright, K. M. Varney, D. J. Weber, L. W. Black, Restriction endonuclease inhibitor IPI* of bacteriophage T4. J Mol Biol 375, 720–734 (2008).

46. E. V. Stabb, E. G. Ruby, “RP4-based plasmids for conjugation between Escherichia coli and members of the Vibrionaceae” in Methods in Enzymology (Academic Press, 2002; https://www.sciencedirect.com/science/article/pii/S0076687902581064)vol. 358 of *Bacte-rial Pathogenesis Part C: Identification, Regulation, and Function of Virulence Factors*, pp. 413–

47. J. Abramson, J. Adler, J. Dunger, R. Evans, T. Green, A. Pritzel, O. Ronneberger, L. Willmore, A. J. Ballard, J. Bambrick, S. W. Bodenstein, D. A. Evans, C.-C. Hung, M. O’Neill, D. Reiman, K. Tunyasuvunakool, Z. Wu, A. Žemgulytė, E. Arvaniti, C. Beattie, O. Bertolli, A. Bridgland, A. Cherepanov, M. Congreve, A. I. Cowen-Rivers, A. Cowie, M. Figurnov, F. B. Fuchs, H. Gladman, R. Jain, Y. A. Khan, C. M. R. Low, K. Perlin, A. Potapenko, P. Savy, S. Singh, A. Stecula, A. Thillaisundaram, C. Tong, S. Yakneen, E. D. Zhong, M. Zielinski, A. Žídek, V. Bapst, P. Kohli, M. Jaderberg, D. Hassabis, J. M. Jumper, Accurate structure prediction of biomolecular interactions with AlphaFold 3. Nature 630, 493–500 (2024).

48. B. Néron, E. Littner, M. Haudiquet, A. Perrin, J. Cury, E. P. C. Rocha, IntegronFinder 2.0: Identification and Analysis of Integrons across Bacteria, with a Focus on Antibiotic Resistance in Klebsiella. Microorganisms 10, 700 (2022).

49. M. Steinegger, J. Söding, MMseqs2 enables sensitive protein sequence searching for the analysis of massive data sets. Nat Biotechnol 35, 1026–1028 (2017).

50. F. Sievers, D. G. Higgins, Clustal Omega for making accurate alignments of many protein sequences. Protein Science 27, 135–145 (2018).

51. S. Khedkar, G. Smyshlyaev, I. Letunic, O. M. Maistrenko, L. P. Coelho, A. Orakov, S. K. Forslund, F. Hildebrand, M. Luetge, T. S. B. Schmidt, O. Barabas, P. Bork, Landscape of mobile genetic elements and their antibiotic resistance cargo in prokaryotic genomes. Nucleic Acids Res 50, 3155–3168 (2022).

52. E. P. Nawrocki, S. R. Eddy, Infernal 1.1: 100-fold faster RNA homology searches. Bioinformatics 29, 2933–2935 (2013).

53. F. Tesson, A. Hervé, E. Mordret, M. Touchon, C. d’Humières, J. Cury, A. Bernheim, Systematic and quantitative view of the antiviral arsenal of prokaryotes. Nat Commun 13, 2561 (2022).

54. M. N. Price, P. S. Dehal, A. P. Arkin, FastTree 2 – Approximately Maximum-Likelihood Trees for Large Alignments. PLOS ONE 5, e9490 (2010).

55. H. Wickham, Ggplot2 (Springer International Publishing, Cham, 2016; http://link.springer.com/10.1007/978-3-319-24277-4)*Use R!*

56. G. Yu, D. K. Smith, H. Zhu, Y. Guan, T. T.-Y. Lam, ggtree: an r package for visualization and annotation of phylogenetic trees with their covariates and other associated data. Methods in Ecology and Evolution 8, 28–36 (2017).

57. A. Kassambara, “ggpubr: ‘ggplot2’ based publication ready plots” (manual, 2023); https://CRAN.R-project.org/package=ggpubr.

58. C. Dawson, “ggprism: a ‘ggplot2’ extension inspired by ‘GraphPad prism’” (manual, 2024); https://csdaw.github.io/ggprism/.

