## Supplementary Materials for "Antiviral defence is a conserved function of diverse DNA glycosylases"

### Materials and Methods

Cloning and expression of glycosylase defence candidates

Candidate glycosylase genes from *Vibrio parahaemolyticus* integrons were synthesized as gene blocks (Twist Bioscience) and cloned into the pVSV105 Vibrio shuttle vector under control of an IPTG-inducible *lac* promoter by In-Fusion cloning (TakaraBio, USA, 638947). All constructs were verified by sequencing. For functional assays, plasmids were transformed into *E. coli* DH5αλpir by heat shock and mobilized into *V. parahaemolyticus* strains by tri-parental conjugation using the helper plasmid pEVS104 (*46*). Where applicable, glycosylase genes were subcloned into pHERD30T for expression in *P. aeruginosa* under the inducible *ara* promoter. All plasmids, primers, and oligonucleotides used in this study can be found in table S10.

Cell culture, phage propagation, and phage plating assays

Phages were propagated on susceptible *E. coli*, *V. parahaemolyticus* or *P. aeruginosa* host strains in liquid culture in lysogeny broth (*E. coli* and *P. aeruginosa,* 10g Tryptone, 5g Yeast Extract, 0.5% NaCl per litre) or lysogeny broth 2% NaCl (LBS, *V. parahaemolyticus*), with the exception of NL2, which was isolated from top-agar following an infection on solid media. To determine phage efficiency of plating, ten-fold serial dilutions of phage lysates were applied to bacterial lawns in LB or LBS top-agar (0.8% agar for *E. coli* and *P. aeruginosa*, 0.4% for *V. parahaemolyticus*) containing the appropriate antibiotics (*E. coli:* chloramphenicol 30 µg/mL, ampicillin 100 µg/mL; *V. parahaemolyticus:* chloramphenicol 2.5 µg/mL; *P. aeruginosa:* gentamycin 50 µg/mL) and inducer (*E. coli* and *V. parahaemolyticus*: 0.1mM IPTG, *P. aeruginosa*: 0.5% arabinose) where necessary. Plating assays were imaged using an EPSON Perfection V850 Pro Photo Scanner in film mode at 800 DPI. Each experiment was performed in biological triplicate; representatives are shown. All bacterial strains used in this study can be found in table S11.

Structural modelling and sequence analysis

AlphaFold3 (*47*) was used to generate structural models of candidate defence proteins in both apo- and DNA-bound states using default parameters. Where indicated, ligands were defined as short dsDNA fragments (5'- GGGGTTTCGCCAGCTGGCGTTTTGGGG-3') containing canonical or modified guanines as indicated, or as single nucleotides. FoldSeek (*29*) analysis was performed to identify structural homologues from the PDB. Putative catalytic residues were inferred by structural alignment with known glycosylases and confirmed by AlphaFold3-predicted binding pockets.

Dag1 protein expression and purification

Dag1 was cloned into p15TV-L with an N-terminal hexahistidine (6xHis) tag to generate 6xHis-Dag1 and the fusion proteins were expressed in *E. coli* BL21(DE3). The proteins were purified using Ni-NTA affinity chromatography and dialyzed into assay buffer (10 mM Tris-HCl pH 6.8, 250 mM NaCl, 0.4mM EDTA, and 5 mM β-mercaptoethanol). Protein purity was confirmed by SDS-PAGE followed by Coomassie staining.

6xHis-Dag1 was subcloned from p15TV-L into VSV105 for expression in *V. parahaemolyticus*, followed by site-directed mutagenesis using Phusion PCR (New England Biolabs, USA,

M0530S) to generate alanine substitution mutants (K118A, D132A, D184A, W188A). Protein expression was induced using 0.1mM IPTG in *V. parahaemolyticus* VP3283-61 at an $OD_{600}$ of 0.8 and allowed to incubate for 3 hours at 30 ℃. Culture density was normalized using $OD_{600}$ and cells were pelleted, followed by lysis in 2x Loading Dye. Protein samples were subjected to SDS-PAGE and western blot. PAGE gels were stained with Coomassie blue. Proteins were transferred to nitrocellulose membranes (0.45 µm, Bio-Rad, USA, 1620115) using the Bio-Rad Trans-Blot SD Semi-Dry Transfer Cell according to manufacturer instructions (Transfer Buffer: 48 mM Tris base, 39 mM glycine, 20 % methanol, 0.2% SDS). Following transfer, membranes were briefly rinsed in TBS and blocked overnight at 4 °C in 5% (w/v) non-fat dry milk (FroggaBio, Canada, SKI400.1) in TBS-T (20 mM Tris-HCl pH 7.4, 150 mM NaCl, 0.1% Tween-20). Membranes were incubated with mouse anti-His primary antibody (BioShop, Canada, TAG001.100; 1:2,000 in TBS-T) for 1 hour at 4 °C with gentle rocking, washed three times for 5 min in TBS-T, followed by incubation with HRP-conjugated rabbit anti-mouse secondary antibody (Cell Signaling Technologies, Canada, 7076S; 1:5,000 in TBS-T) for 1 h at 4 °C. Western blots were developed using the Clarity Western ECL Substrate kit (Bio-Rad, USA, 1705060).

Phage genome extraction, gDNA restriction analysis and *in vitro* DNA degradation

Following treatment with DNaseI (Thermo Scientific, USA, EN0521) and RNase A (Thermo Scientific, USA, EN0531) to remove host nucleic acids leftover from cell lysis, phage gDNA was purified using standard phenol/chloroform DNA extraction protocols (Invitrogen, USA, 15593031). Phage genomes were subjected to restriction analysis using EcoRV, HaeIII, NdeI or SspI (New England Biolabs, USA) using manufacturer protocols.

To test the *in vitro* activity of Dag1 against phage gDNA, variable concentrations of purified 6xHis-Dag1 were incubated with 200 ng of phage genomic DNA at 37 °C for 30 min in assay buffer as above. Reactions were quenched with loading dye and run on 1% agarose gels stained with SYBR-Safe (Invitrogen, USA, S33102; 100V/Room Temperature). The same DNA degradation assays were also visualized on 1% agarose gels run at 20V/4°C to avoid degradation of abasic DNA by high voltage and heat.

Phage genome modification analysis

Phage genomic analysis was performed using Domainator using custom HMMs against known DNA modification enzymes (*24*) and visualized with Clinker (https://github.com/gamcil/clinker). To analyze nucleoside content of phage gDNA, genomic DNA from phages HK243, HK243 $DpdA^{G217E}$, VP1, and NL2 was extracted as above and analyzed by HPLC-MS as previously described (*35*). Canonical and modified nucleosides (e.g., $dPreQ_0$, 7-deazaguanine) were identified based on retention time compared to nucleoside standards and their associated mass spectra. Restriction digestion assays were performed with Type II enzymes predicted to cut unmodified phage DNA. Digestion products were analyzed on 1% agarose gels stained with SYBR-Safe (Invitrogen, USA, S33102).

Selection and analysis of phage escaper mutants

Phage HK243 was plated on *E. coli* DH5αλpir expressing Dag1 from a plasmid. Following overnight incubation, resistant plaques appeared on Dag1-expressing lawns. Individual escaper phages were isolated and purified through two rounds of plaque purification in the presence of Dag1. Phage gDNA was extracted from three escape mutants and subjected to whole-genome sequencing using Oxford Nanopore Technologies. gDNA was analyzed using snippy (https://github.com/tseemann/snippy). All three escaper phages carried identical G217E mutations in *dpdA*, a gene implicated in the incorporation of 7-deazaguanine modifications.

Vibrionacea integron sequence database generation

Vibrionacea genomes were downloaded from NCBI on January 27th 2025, using NCBI datasets (taxid: 641, --assembly-level chromosome,complete). Only complete or nearly complete genomes were collected to ensure minimal contig breaks within the integron. Genomes were deduplicated manually, preferentially including RefSeq accessions. Genbank accessions were included where no RefSeq annotated genome was available.

IntegronFinder2.0.2 (*48*) was used to identify complete sedentary chromosomal integrons (using –local_max). Integron regions were summarized using a custom python script, and protein sequences annotated within these regions were collected from NCBI using a custom python script. Amino acid sequences were clustered at 80% pairwise coverage and 30% sequence identity by mmseqs2 (v13.45111, easy-cluster –cov-mode 0) (*49*).

ESM3 structure database generation and structural homologue searches

Protein structures were generated for the representativeintegron-encoded proteins using ESM3 (*28*) and the ESM3-open model available on HuggingFace (esm3-sm-open-v1). Structures were folded using ESM3's round-trip design approach to refine the structure output, using a custom python script. Following protein structure generation, FoldSeek was used to generate a structural database used below (v10.941cd33, createdb, default parameters).

FoldSeek was used to determine structurally similar protein homologue using easy-search mode against the previously generated ESM3 FoldSeek database. FoldSeek results were filtered for pairwise query/target coverage greater than 60%, an E-value of less than 2, and calculated RMSD over aligned portions of the structures less than 15. Large E-values were permitted to ensure maximum diversity in the structures obtained. Structural homologues were then manually inspected through alignment using ChimeraX to ensure alignment in the core glycosylase HhH-GPD motif, and those with poor alignment were removed. Brig1 homologues were identified using the same approach. Following identification, homologue were subjected to Clustal Omega (*50*) alignments to determine protein sequence identity, AlphaFold3 (*47*) to determine structural confidences and predicted structure, and DALI structural comparison analysis (*30*) using the online servers for each tool.

Homologue, taxonomy, and genomic context analysis of structural homologues

Sequence identity matrices for the identified homologues were performed following sequence alignment with Clustal Omega (v1.2.4, default settings) (*50*). Each homologue was subjected to PSI-BLAST analysis against the NCBI nr database (-num_iterations 10 -evalue 1e-3). Identified PSI-BLAST matches (filtering for >30% sequence identity and >70% alignment coverage) were then used to identify taxonomy and distribution of their associated genomes using NCBI's Identical Protein Groups (IPG) database, using a custom python script. The IPG results were made non-redundant by removing identical assemblies from the dataset.

To analyze the context surrounding each identified homologue, each PSI-BLAST match was used to query the IPG database, followed by extraction of associated genomes (up to a maximum of 100 genomes per accession) and subsequent deduplication of identical genomes. 20kb regions surrounding the homologue were subsequently extracted and subjected to further analysis. Only genomic context regions at least 20kb in size were included to ensure sufficient context in relation to whole genomic fragments to provide an accurate analysis of G/C content differences. G/C content was calculated for both extracted regions, and protein annotations within the 20kb region were queried by keyword, by HMM using supervised HMMs generated by Khedkar *et al.* (*51*), and by cmsearch using IntegronFinder2 *attC* queries (*48*, *52*) for common mobile-genetic element associated features (*e.g.* recombinases, integrases, transposases, and integron *attC* sites). G/C Content difference was calculated as the difference between genomic G/C content and the G/C content of the 20kb region. DefenseFinder was used to identify known antiphage defences in these regions (*53*).

Phylogenetic Analysis and Tree Construction

Sequences to be compared were downloaded from NCBI using Batch Entrez and then aligned using ClustalO (v1.2.4, default settings) (*50*). FastTree (v2.2) (*54*) was used to construct the tree using default settings and R packages 'ggplot2' and 'ggtree' were used to visualize and annotate the tree.

Data visualization and statistical analysis

Unless otherwise specified, plots were generated in R using ggplot2 (*55*), ggtree (*56*), ggpubr (*57*), and ggPrism packages (*58*). Structural models were visualized and analyzed in ChimeraX.

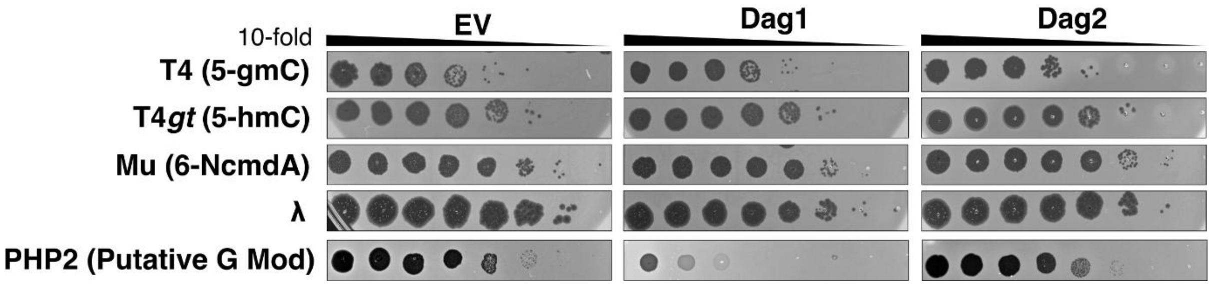


**Fig. S1.**

**Additional phage plating data for Dag1 and Dag2.** Phage plating assays of modified cytosine phages T4 (5-gmC), T4*gt* (5-hmC), Mu (6-NcmdA), λ (unmodified), and PHP2 (unconfirmed G-modification) on cells expressing Dag1 and Dag2. Images are representative of two biological replicates.

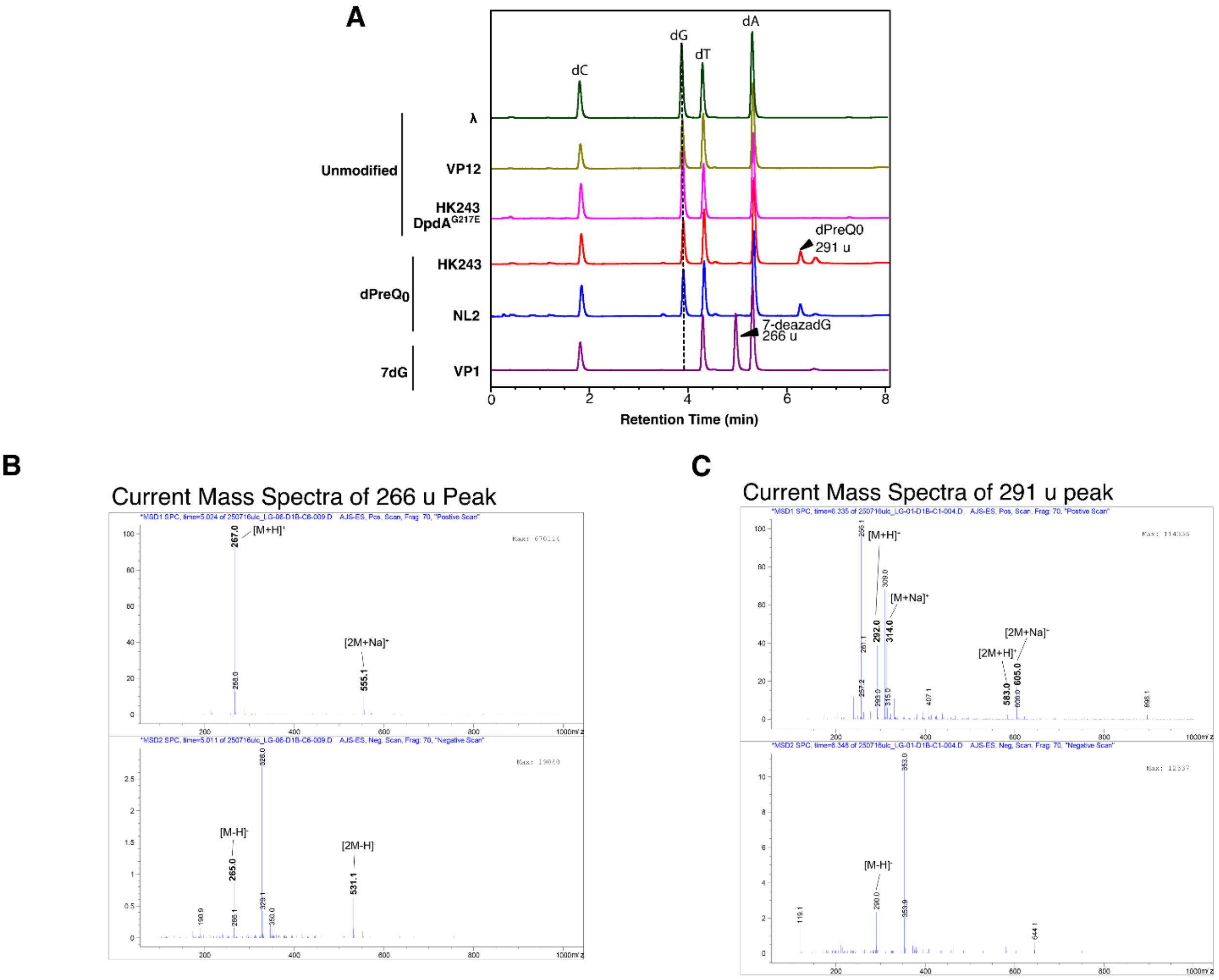


**Fig. S2.**

**High-performance liquid chromatography and mass spectrometry data for a select panel of phage genomes.** (**A**) HPLC traces for phages lambda, VP12, HK243, NL2, VP1 and HK243 DpdA$^{G217E}$. The dG peak is indicated across the traces by a dotted line. Peaks corresponding to modified guanosines are labelled. Mass spectra evidence for the 7-deazaguanine (266 u) (**B**) and dPreQ0 (291 u) (**C**) peaks in (**A**).

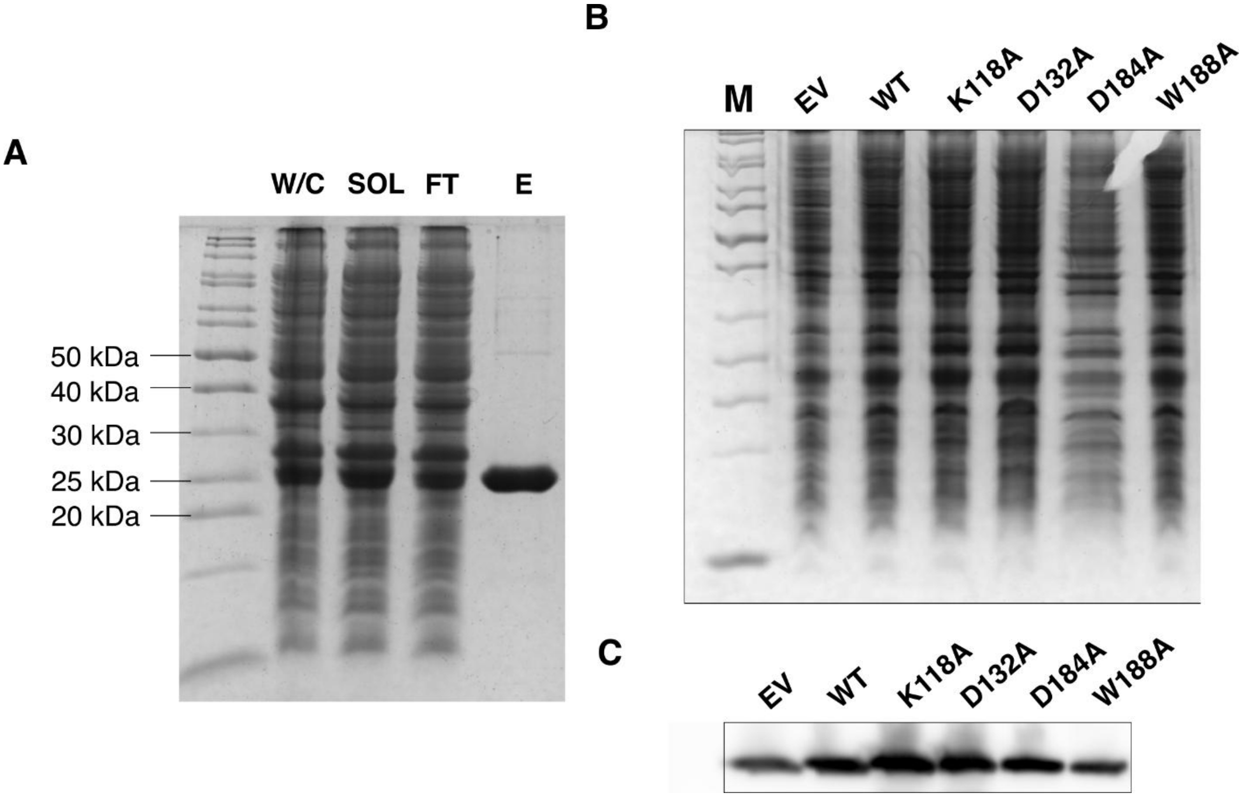


**Fig. S3.**

**Site-directed mutagenesis of key Dag1 active site residues identifies those necessary for function**. (**A**) 6xHisDag1 was expressed in BL21(DE3) *E. coli* and whole cell (W/C), soluble (SOL), flow-through (FT) and elution (E) fractions were subjected to SDS-PAGE stained with Coomassie blue. (**B**) Coomassie stained SDS-PAGE gel of whole cells lysates of *V. parahaemolyticus* 3283-61 cells expressing each of the Dag1 alanine-substitution mutants. (**C**) Anti-6xHis immunoblot against the same samples shown in (**B**).

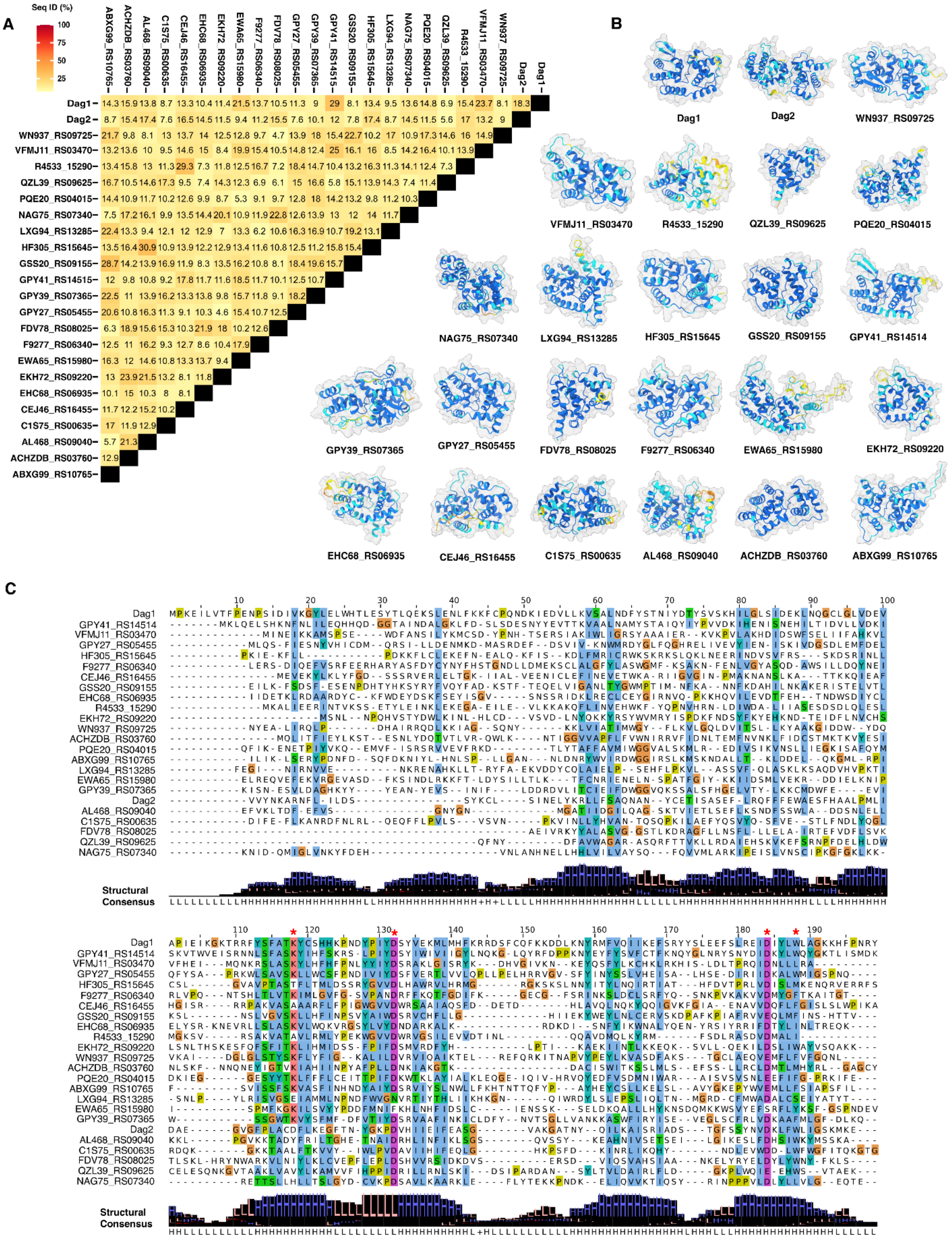


**Fig. S4.**

**Dag1 and Dag2 structural homologues are abundant in Vibrionacea integrons.** (A) Dag1 and Dag2 structural homologues were identified by FoldSeek and compared by multiple sequence alignment using

Clustal Omega (*29*, *50*). ClustalO percent identity matrices were used to generate the plot shown using R and ggplot2. Locus tags are used where no defence phenotype was described, and Dag identifiers are used for those which have defence phenotypes according to this study. (**B**) AlphaFold3 was used to generate structural predictions of each homologue and are shown. Structures are coloured by pLDDT at each residue (dark blue, pLDDT >90; light blue, 90 > pLDDT > 70; yellow, low 70 > pLDDT > 50; orange, very low pLDDT<50). (**C**) DALI generated structural alignments, without gaps, are shown. Sequence alignments were visualized using JalView and are coloured by Clustal colouring. The structural consensus is shown underneath each position. The residues involved in catalysis and substrate recognition are indicated by red asterisks.

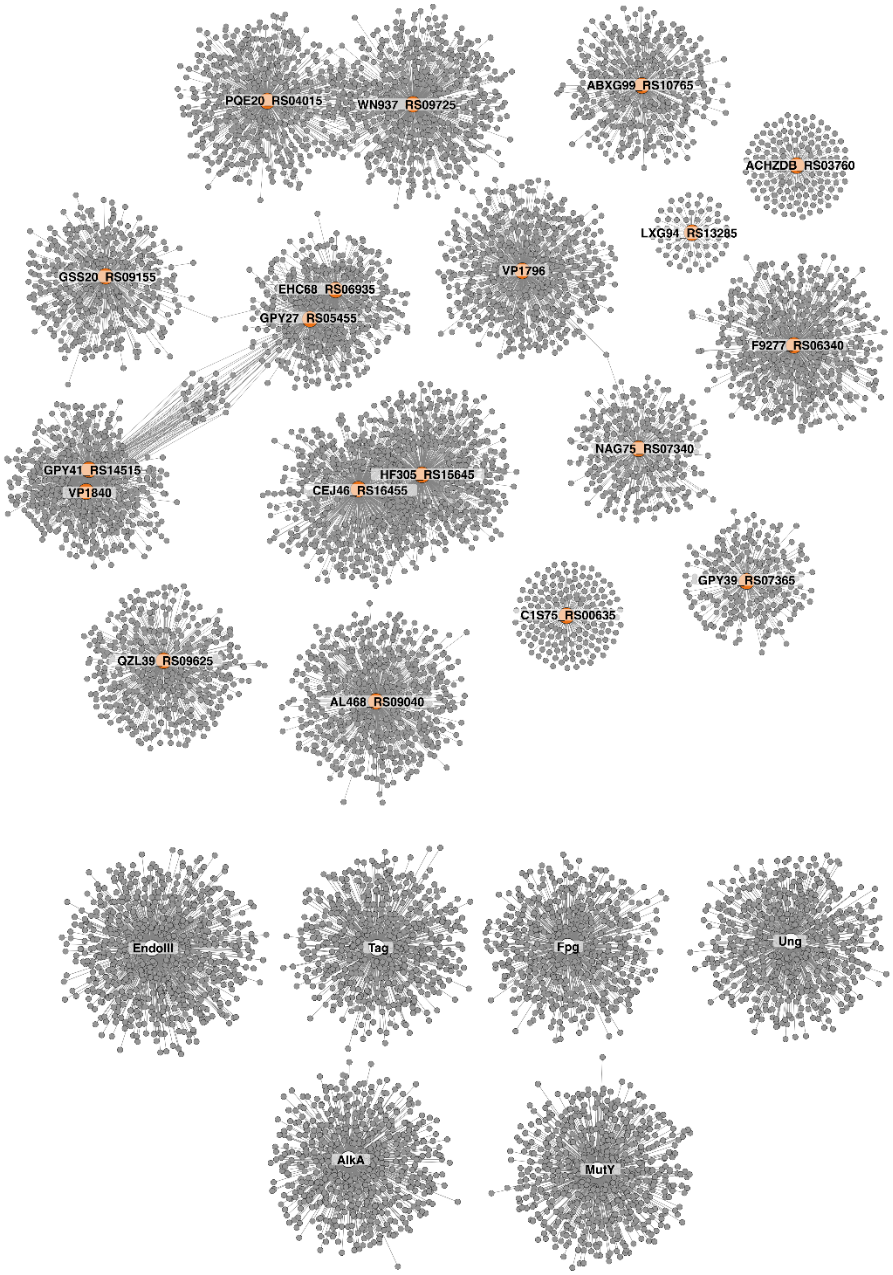


**Fig. S5.**

**PSI-BLAST network analysis of Dag homologues and *V. parahaemolyticus* housekeeping glycosylases.** PSI-BLAST was performed to 10 iterations (or convergence) using an E-value inclusion criteria of 0.1, selecting only homologues with greater than 70% pairwise coverage (all other settings were default). Query and subjects were linked by network analysis and visualized using Cytoscape3. Grey dots indicate subject proteins matching the Dag queries by PSI-BLAST. Shared nodes indicate overlapping PSI-BLAST matches. Orange queries have antiphage function, while white queries do not.

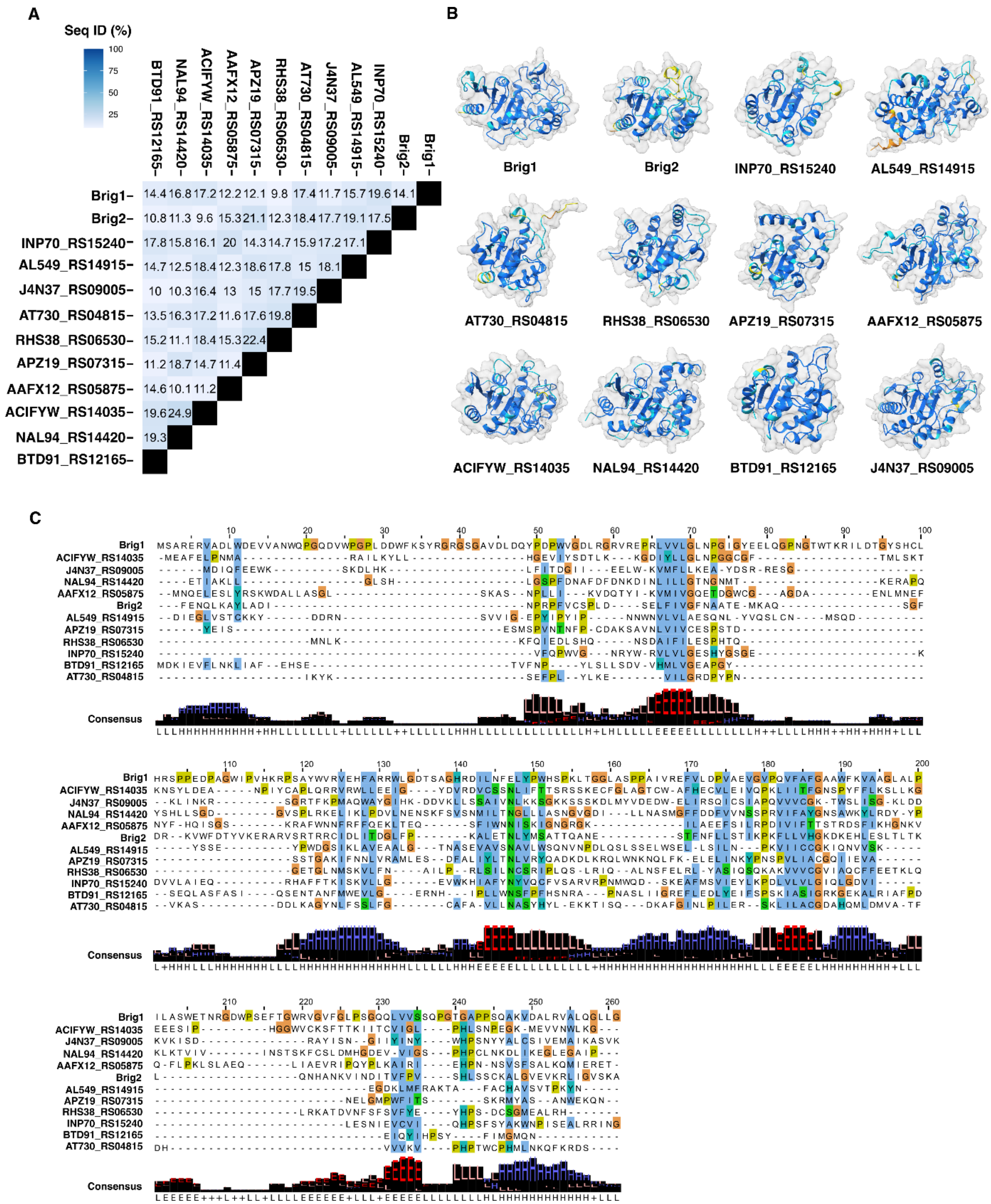


**Fig. S6**

**Brig1 homologues are also abundant in the Vibrionacea integrons.** (**A**) Brig1 structural homologues were identified by FoldSeek and compared by multiple sequence alignment using ClustalO (*29*, *50*). ClustalO percent identity matrices were used to generate the plot shown. Locus tags are used where no defence phenotype was described, and additional Brig identifiers are used for those which have defence phenotypes according to this study. (**B**) AlphaFold3 was used to generate structural predictions of each homologue and are shown. Structures are coloured by pLDDT at each residue (dark blue, pLDDT >90; light blue, 90 > pLDDT > 70; yellow, low 70 > pLDDT > 50; orange, very low pLDDT<50). (**C**) DALI generated structural alignments, without gaps, are shown. Sequence alignments were visualized using JalView and are coloured by Clustal colouring. The structural consensus is shown underneath each position.

**Table S1.** Summary table of DNA glycosylases found in Bacteria and Archaea

**Table S2.** List of phages used in this study

**Table S3.** Representative proteins identified using Dag1 and Dag2 as FoldSeek queries against integron protein database

**Table S4.** PSI-BLAST results for the representative proteins from Table S3 with antiphage function and assigned Dag nomenclature

**Table S5.** Taxonomic assignment to homologues of functional Dag and Brig2 family members

**Table S6.** Housekeeping glycosylases identified in representative *Vibrio* species genomes

**Table S7.** DefenseFinder Results for 20kb Dag family member genomic contexts

**Table S8.** Genomic context of Dag family members, housekeeping glycosylases, and Brig2

**Table S9.** Representative proteins identified using Brig1 as a FoldSeek queries against integron protein database

**Table S10.** List of oligonucleotides used in this study

**Table S11.** List of bacterial strains used in this study
